# Decomposing drivers in avian insectivory: large-scale effects of climate, habitat and bird diversity

**DOI:** 10.1101/2023.01.19.524212

**Authors:** Laura Schillé, Elena Valdés-Correcher, Frédéric Archaux, Flavius Bălăcenoiu, Mona Chor Bjørn, Michal Bogdziewicz, Thomas Boivin, Manuela Branco, Thomas Damestoy, Maarten de Groot, Jovan Dobrosavljević, Mihai-Leonard Duduman, Anne-Maïmiti Dulaurent, Samantha Green, Jan Grünwald, Csaba Béla Eötvös, Maria Faticov, Pilar Fernandez-Conradi, Elisabeth Flury, David Funosas, Andrea Galmán, Martin M. Gossner, Sofia Gripenberg, Lucian Grosu, Jonas Hagge, Arndt Hampe, Deborah Harvey, Rick Houston, Rita Isenmann, Andreja Kavčič, Mikhail V. Kozlov, Vojtech Lanta, Bénédicte Le Tilly, Carlos Lopez Vaamonde, Soumen Mallick, Elina Mäntylä, Anders Mårell, Slobodan Milanović, Márton Molnár, Xoaquín Moreira, Valentin Moser, Anna Mrazova, Dmitrii L. Musolin, Thomas Perot, Andrea Piotti, Anna V. Popova, Andreas Prinzing, Ludmila Pukinskaya, Aurélien Sallé, Katerina Sam, Nickolay V. Sedikhin, Tanja Shabarova, Ayco J. M. Tack, Rebecca Thomas, Karthik Thrikkadeeri, Dragoș Toma, Grete Vaicaityte, Inge van Halder, Zulema Varela, Luc Barbaro, Bastien Castagneyrol

**Author notes:** Luc Barbaro and Bastien Castagneyrol share last co-authorship.

## Abstract

**Aim:** Climate is a major driver of large scale variability in biodiversity, as a likely result of more intense biotic interactions under warmer conditions. This idea fuelled decades of research on plant-herbivore interactions, but much less is known about higher-level trophic interactions. We addressed this research gap by characterizing both bird diversity and avian predation along a climatic gradient at the European scale.

**Location:** Europe.

**Taxon:** Insectivorous birds and pedunculate oaks.

**Methods:** We deployed plasticine caterpillars in 138 oak trees in 47 sites along a 19° latitudinal gradient in Europe to quantify bird insectivory through predation attempts. In addition, we used passive acoustic monitoring to (i) characterize the acoustic diversity of surrounding soundscapes; (ii) approximate bird abundance and activity through passive acoustic recordings and (iii) infer both taxonomic and functional diversity of insectivorous birds from recordings.

**Results:** The functional diversity of insectivorous birds increased with warmer climates. Bird predation increased with forest cover and bird acoustic activity but decreased with mean annual temperature and functional richness of insectivorous birds. Contrary to our predictions, climatic clines in bird predation attempts were not directly mediated by changes in insectivorous bird diversity or acoustic activity, but climate and habitat still had independent effects on predation attempts.

**Main conclusions:** Our study supports the hypothesis of an increase in the diversity of insectivorous birds towards warmer climates, but refutes the idea that an increase in diversity would lead to more predation and advocates for better accounting for activity and abundance of insectivorous birds when studying the large-scale variation in insect-tree interactions.

## Résumé

**Objectif:** Le climat est l’un des principaux facteur structurant de la variabilité à grande échelle de la biodiversité, possiblement en raison d’interactions biotiques plus intenses dans des conditions de température plus élevées. Cette idée a alimenté des décennies de recherche sur les interactions plantes-herbivores, mais on en sait beaucoup moins sur les interactions impliquant les niveaux trophiques supérieurs. Nous avons comblé cette lacune en caractérisant à la fois la diversité des oiseaux et leur activité de prédation le long d’un gradient climatique à l’échelle européenne.

**Localisation:** Europe.

**Taxon:** Oiseaux insectivores et chênes pédonculés.

**Méthodes:** Nous avons déployé des leurres en pâte à modeler mimant des chenilles sur 138 chênes dans 47 sites le long d’un gradient latitudinal de 19° en Europe pour quantifier l’insectivorie avienne par le biais de tentatives de prédation. De plus, nous avons utilisé la surveillance acoustique passive pour (i) caractériser la diversité acoustique des paysages sonores environnants ; (ii) estimer l’abondance et l’activité des oiseaux à travers des enregistrements acoustiques passifs et (iii) déduire à la fois la diversité taxonomique et fonctionnelle des oiseaux insectivores à partir des enregistrements.

**Résultats:** Nous avons montré une augmentation de la diversité fonctionnelle des oiseaux insectivores avec la température moyenne. La prédation avienne augmentait avec la couverture forestière et l’activité acoustique des oiseaux, mais diminuait avec la température annuelle moyenne et la richesse fonctionnelle des oiseaux insectivores. Contrairement à nos prédictions, la variation de la diversité des oiseaux n’était pas le lien mécaniste entre le climat et la variation des tentatives de prédation sur les leurres, laquelle était directement influencée par le climat et la couverture forestière.

**Conclusions principales:** Notre étude confirme l’hypothèse d’une augmentation de la diversité des oiseaux insectivores vers des climats plus chauds, mais ne corrobore pas l’idée qu’une augmentation de la diversité conduirait à davantage de predation. Elle plaide en faveur d’une meilleure prise en compte de l’activité et de l’abondance des oiseaux insectivores lors de l’étude de la variation à grande échelle des interactions entre insectes et arbres.

## Introduction

Climate is a key driver of biotic interactions (Dobzhansky, 1950). A long held view in ecology posits that warmer and more stable climatic conditions intensify biotic interactions and accelerates speciation (MacArthur, 1984; Schemske, Mittelbach, Cornell, Sobel & Roy, 2009), which should result in large scale positive correlations between biodiversity and biotic interactions. However appealing this idea is, the generality of large-scale climatic clines in biodiversity and biotic interactions as well as the underlying causal links are still widely debated. Yet, insights into the controversy have been dominated by studies on plant-insect interactions (Anstett, Chen & Johnson, 2016; Kozlov, Lanta, Zverev & Zvereva, 2015). Biotic interactions involving higher trophic levels received much less attention. Yet, insectivorous birds are among the predators contributing the most to the control of insect herbivores in terrestrial ecosystems (van Bael et al., 2008; Sam, Jorge, Koane, Amick & Sivault, 2023; Sekercioglu, 2006) and therefore have consequences on both the assembly of ecological communities and the functioning of ecosystems. The omission of predation in theories linking large-scale variability in climate with biodiversity therefore represents a critical gap in knowledge that needs to be addressed.

Bird communities are highly responsive to climate, at both regional and continental scale. There is a large body of literature demonstrating that several dimensions of bird diversity vary with climate, including bird abundance, species richness, phylogenetic or functional diversity (Blackburn & Gaston, 1996; Symonds Christidis & Johnson, 2006; Willig, Kaufman & Stevens, 2003). A well substantiated explanation is that niche opportunities increase with increasing habitat heterogeneity under milder climatic conditions, which increases species coexistence and ultimately species richness through functional complementarity (Hawkins, Diniz-Filho, Jaramillo & Soeller, 2006). The biodiversity and ecosystem relationship theory predicts that both abundance and diversity of birds are crucial predictors of the top-down control they exert upon insect prey (Bael et al., 2008; Nell, Abdala-Roberts, Parra-Tabla & Mooney, 2018; Otto, Berlow, Rank, Smiley & Brose, 2008; Sinclair, Mduma & Brashares, 2003). Numerous studies supported this theory and demonstrated that bird functional diversity in particular --- that is the diversity, distribution and complementarity of predator traits involved in predation --- is a good predictor of predation (Barbaro, Giffard, Charbonnier, van Halder & Brockerhoff, 2014; Greenop, Woodcock, Wilby, Cook & Pywell, 2018; Philpott et al., 2009). It follows that variation in bird diversity along climatic gradients should be mirrored by consistent variation in avian predation rates.

Local factors can however alter macroecological patterns (Ikin et al., 2014; Kissling, Sekercioglu & Jetz, 2012), by filtering the regional species pool (De la Mora, García-Ballinas & Philpott, 2015; Kleijn, Rundlöf, Scheper, Smith & Tscharntke, 2011) and by influencing the behavior of organisms. The diversity and composition of bird communities heavily depends on local factors that provide niches and food opportunities (Charbonnier et al., 2016). In this respect, multiscale forest cover proved to be a particularly good predictor of composition of birds communities at different spatial scales, as bird foraging activity is ultimately determined by vertical and horizontal habitat heterogeneity, which influences both where prey can be found and caught, and where foraging birds can breed and hide from predators (Vickery & Arlettaz, 2012). Thus, modeling the response of bird communities to large-scale bioclimatic drivers as well as their role as predators would benefit from using a combination of habitat variables and biotic predictors (Barbaro et al., 2019; Speakman et al., 2000). However, cross-continental studies exploring the relationship between large scale climatic gradients and the strength of biotic interactions generally ignore local factors, which may partly explain inconsistencies in their findings (but see Just, Dale, Long & Frank, 2019).

A major challenge to analyze climatic clines in biotic interactions consists in simultaneously characterizing changes in predator biodiversity and experimentally assessing the strength of predation, while considering the effect of contrasting habitats. However, the recent development of passive acoustic monitoring provides a standardized, low-cost and non-invasive approach for ecological studies and biodiversity monitoring (Gibb, Browning, Glover-Kapfer, Jones & Börger, 2019). The acoustic monitoring of a given habitat primarily allows the delayed identification of bird species over large gradients with no need for distributed expertise across study sites. The quantification of bird abundance through passive acoustic monitoring remains a technical challenge, but the calculation of certain acoustic indices based on the physical characteristics of the recorded sounds provides relevant proxies to this end (Gasc et al., 2013; Sueur, Farina, Gasc, Pieretti & Pavoine, 2014). Should such indices consistently correlate with macro-scales biotic interactions, ecoacoustics would be a promising complementary approach to existing methods in macroecology and in functional ecology.

Here, we addressed the hypothesis of continental north-south clines on insectivorous bird community diversity and their predation function, while controlling for local factors throughout the European distribution range of the pedunculate oak (*Quercus robur* L., 1753), a major forest tree species. Specifically, we predict the following (Fig. 1): (i) bird diversity (including bird acoustic diversity, insectivorous bird species richness and functional diversity) and predation attempts increase with warmer climates; (ii) bird predation attempts increase with bird acoustic activity, species richness and greater functional diversity of insectivorous birds; (iii) bird diversity, acoustic activity and bird predation attempts increase with increasing forest cover at both local (neighborhood) and larger spatial scales; (iv) large-scale variability in bird predation attempts is driven by local changes in the diversity and acoustic activity of birds. To test these predictions, we quantified bird predation attempts on plasticine caterpillars and estimated bird species richness, functional diversity and acoustic activity through simultaneous passive acoustic monitoring. We eventually tested the respective responses of these variables and their relationships at the pan-European scale.

**Figure 1:**
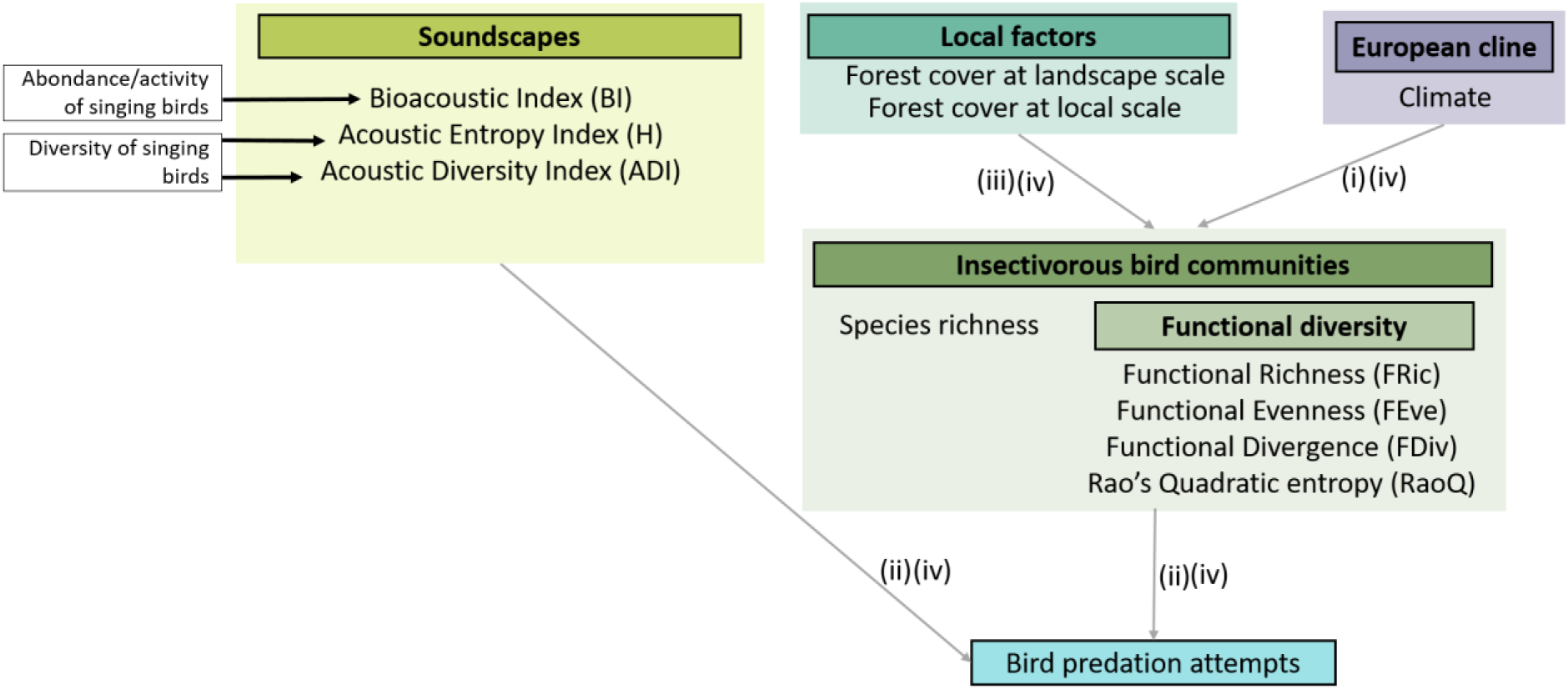
Conceptual diagram of the predictions of this study and the relationships already established in the literature. Boxed elements written in bold correspond to the main categories of variables tested, they are not variables as such. Variables used in models are shown in regular font. Where several variables described the same category (e.g. BI, ADI, H, all describing acoustic indices), we used multi-model comparisons to identify the best variable. Items framed in black on a white background represent untested variables. Black arrows indicate relationships well supported by the literature (see Gasc et al., 2018; and Fig.2 Sánchez-Giraldo, Correa Ayram & Daza, 2021). Our specific predictions are represented with grey arrows, solid and dashed lines representing positive and negative (predicted) relationships. Numbers refer to predictions as stated in the main text.

## Materials and methods

### Study area

We focused on the pedunculate oak, *Quercus robur,* which is one of the keystone deciduous tree species in temperate European forests, where it is of high ecological, economic and symbolic importance (Eaton, Caudullo, Oliveira & de Rigo, 2016). The species occurs from central Spain (39°N) to southern Fennoscandia (62°N) and thus experiences a huge gradient of climatic conditions (Petit et al., 2002). A widely diverse community of specialist and generalist herbivorous insects is associated with this species throughout its distributional range (Southwood, Wint, Kennedy & Greenwood, 2005).

Between May and July 2021, we studied 138 trees in 47 sites across 17 European countries covering most of the pedunculate oak geographic range (Fig. 2). The sites were chosen with the minimal constraint of being located in a wooded area of at least 1 ha (Valdés-Correcher et al., 2021). We randomly selected three mature oaks per site, with the exception of six sites (three sites with one tree, one site with two trees and two sites with five trees, see Table S1.1 in Appendix S1 in Supporting Information).

**Figure 2:**
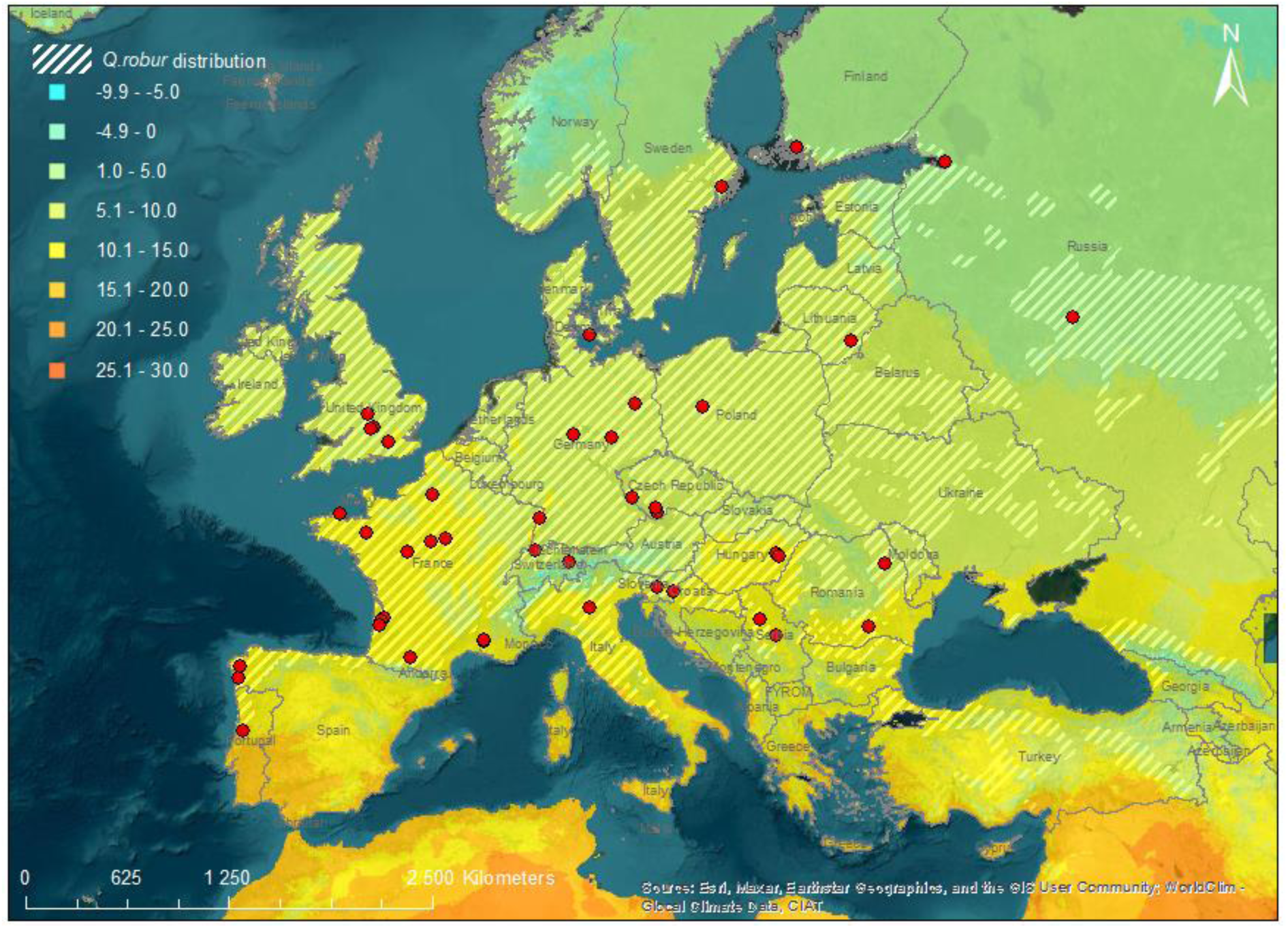
Locations of the 47 sites sampled in spring 2021. Average annual temperature (color scale) according to WorldClim (Hijmans, Cameron, Parra, Jones & Jarvis, 2005) and Quercus robur distribution range are indicated.

### Bird predation attempts

We measured bird predation attempts in the field by exposing a total of 40 plasticine caterpillars (20 plasticine caterpillars twice) on each individual oak. We made plasticine caterpillars of green plasticine, mimicking common lepidopteran larvae (3 cm long, 0.5 cm diameter, see Low, Sam, McArthur, Posa & Hochuli, 2014). We secured them on twigs with a 0.3 mm metallic wire. We attached five plasticine caterpillars to each of four branches facing opposite directions (i.e., 20 caterpillars per tree) at about 2 m from the ground.

We installed the plasticine caterpillars six weeks after budburst in each study area, thus synchronizing the study with local oak phenology. We removed the plasticine caterpillars after 15 days and installed another set of 20 artificial caterpillars per tree for another 15 days. At the end of each exposure period (which varied from 10 to 20 (mean ± SD: 14.5 ± 1.23) days due to weather conditions, we carefully removed the plasticine caterpillars from branches, placed them into plastic vials and shipped them to the project coordinator. Plasticine caterpillars from six sites were either lost or altered during shipping, preventing the extraction of relevant data.

A single trained observer (EVC) screened the surface of plasticine caterpillars with a magnifying lens to search for the presence of bill marks on clay surface (Low et al., 2014). As we were ultimately interested in linking bird diversity with bird predation attempts, we did not consider marks left by arthropods and mammals.

We defined *bird predation attempts index* as *p / d*, where *p* is the proportion of plasticine caterpillars with at least one sign of attempted predation by birds and *d* is the number of days plasticine caterpillars were exposed to predators in the field. We only considered as attacked those caterpillars that we retrieved; missing caterpillars were not accounted for in the calculation of *p*. We calculated bird predation attempts for each tree and survey period separately. Because other variables were defined at site level (see below), we averaged bird predation attempts across trees and surveys in each site (total: *n* = 41).

To assess the effect of temperature independently of other variables that could vary with latitude, we also calculated a second bird predation attempts index by standardizing the predation attempts by daylight duration in every site. We ran the same statistical models as for the non-standardized bird predation attempts. The outcomes remained qualitatively the same and the results of this analysis are presented in Table S2.2 in Appendix S2.

### Acoustic monitoring and related variables

We used passive acoustic monitoring to characterize the species and functional diversity of bird communities associated with oaks, as well as to serve as a proxy of the abundance and diversity of vocalizing birds (Fig. 2). In each site, we randomly chose one oak among those used to measure bird predation rates in which we installed an AudioMoth device (Hill et al., 2018) to record audible sounds for 30 min every hour. Automated recording started the day we installed the first set of 20 plasticine caterpillars in trees and lasted until batteries stopped providing enough energy. The recording settings were the following: Recording period: *00.00-24.00 (UTC); Sample rate: 48 kHZ; Gain: Medium; Sleep duration: 1800 s, Recording duration: 1800 s*.

In all 47 sites, Audiomoths were active on average (± S.D.) for 9 ± 3 days (range: 1-24), which corresponded to 5920 h of recordings in total and from 70 to 335 (246 ± 65) 30 min continuous acoustic samples per site. When Audiomoths ran out of battery, the recordings lasted less than 30 min (between 1 and 56 recordings per site were affected).

### Acoustic diversity indices as proxies of bird diversity and activity

We processed acoustic samples with functions in the “soundecology” v.1.3.3 (Villanueva-Rivera & Pijanowski, 2018) and “seewave” v. 2.1.8 (Sueur, Aubin & Simonis, 2008) libraries in the R environment version 4.1.2 (R Core Team, 2020), and a wrap-up function made available by A. Gasc in GitHub (https://github.com/agasc/Soundscape-analysis-with-R). We first divided every acoustic sample (regardless of its length) into non-overlapping 1 min samples.

Acoustic indices capture various dimensions of the soundscape but are not expected to fully reflect any bird biodiversity-related variable. However, several studies have shown that some of them are positively related to the abundance or diversity of vocalizing species (for more details, see Sánchez-Giraldo, Correa Ayram & Daza, 2021, Fig.2 and Gasc et al., 2018), although the strength of this relationships is still poorly understood. We have therefore chosen to consider only those specific indices and we used multi-model statistical inferences to identify those that were the most strongly linked with the response variables of interest (see below).

We calculated the following three acoustic diversity indices for each 1 min sample: the Acoustic Diversity Index (ADI) and the Total Acoustic Entropy (H) which are both based on Shannon diversity index and are therefore close to a proxy for bird diversity (Sueur, Pavoine, et al., 2008; Villanueva-Rivera, Pijanowski, Doucette & Pekin, 2011), and the Bioacoustic Index (BI) which is positively related to bird vocal activity and the occupancy of acoustic signal frequency bands (Boelman, Asner, Hart & Martin, 2007; Gasc et al., 2018). We calculated the median of each acoustic index per day and then averaged median values across days for each site separately. We proceeded like this because 24 h cycles summarize the acoustic activity and account for all possible sounds of a given day. Furthermore, other studies have previously shown that median values of acoustic indices for a given day are more representative than mean values of the acoustic activity because they are less sensitive to extreme values (Barbaro et al., 2022; Dröge et al., 2021). This procedure resulted in one single value of each acoustic diversity index per site.

### Bird species richness and functional diversity

We used acoustic samples to identify birds based on their vocalizations (songs and calls) at the species level, from which we further computed functional diversity indices (Fig. 3).

#### Data processing

For each site, we subsampled the 30 min samples corresponding to the songbird morning chorus (*i.e.,* the period of maximum singing activity), which incidentally also corresponds to the time of the day when anthropic sounds were of the lowest intensity. Specifically, we selected sounds recorded within a period running from 30 min before sunrise to 3 h 30 min after sunrise. We then split each 30 min sample into up to three 10 min sequences, from which we only retained those recorded on Tuesday, Thursday, Saturday, and Sunday. We chose these days on purpose to balance the differences in anthropogenic noises between working days and weekends. For each sound sample, we displayed the corresponding spectrogram with the “seewave” library in the R environment (Sueur, Aubin & Simonis, 2008). We visually sorted sound samples thanks to spectrograms and discarded samples with noise from anthropogenic sources, rain, or wind, which can be recognized as very low frequency noise on the spectrogram. We also discarded samples with noise of very high frequency corresponding to cicada chirps. We then randomly selected one sound sample per site and per day, with the exception of four sites for which the four samples only covered two to three days. In total, we selected 188 samples of 10 min (i.e., 4 samples per site).

**Figure 3:**
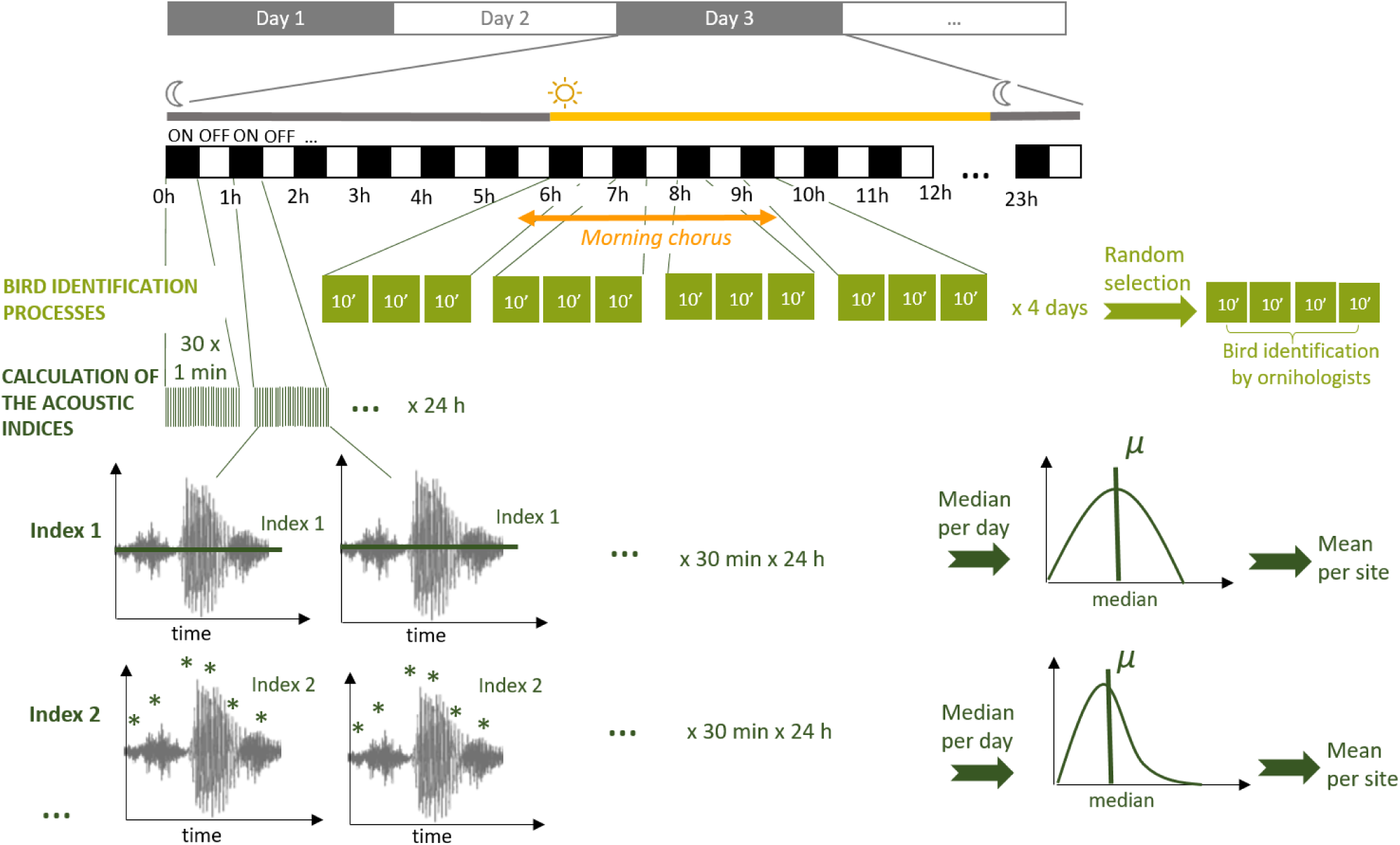
Methodological pathway used to identify bird species (in light green) and calculate acoustic indices (in dark green) from automated recordings (see text for details)

#### Bird species identification

We distributed the samples among 21 expert ornithologists. Each expert performed aural bird species identifications from 4 (one site) to 52 samples (13 sites), primarily from her/his region of residence, for auditory acoustic detection of bird species. We established a presence/absence Site × Species matrix, from which we calculated species richness and functional diversity. It is important to note that, there is no possibility to determine the direction and distance at which birds are singing from audio recordings when using a single device for a given site. As a result, there is no standard method for determining whether or not two vocalizations of the same species at two different times come from one single individual or more, which prevents an accurate estimate of bird abundance. However, experienced ornithologists involved in this study consider that, given the territoriality of birds and the range of the recorders, it is unlikely that they recorded the vocalizations of several individuals of the same species. It therefore seems reasonable to assume that among-site differences in bird species richness were also representative of among-site differences in bird abundance.

#### Functional diversity

We defined 25 bird species as candidate insectivores for attacking plasticine caterpillars (Table S3.3 in Appendix S3) with those bird species meeting the following criteria: be insectivorous during the breeding season or likely to feed their offspring with insects, forage primarily in forested habitats, and are likely to use substrates such as lower branches or lower leaves of trees where caterpillars were attached to find their prey (Barbaro et al., 2021; Brambilla & Gatti, 2022). We calculated the functional diversity of these candidate insectivores by combining morphological, reproductive, behavioral and acoustic traits.

With the exception of acoustic traits, we extracted functional traits from different published sources, listed in Table S3.4 in Appendix S4. Specifically, we used three continuous traits: *body mass*, *mean clutch size* and *bill culmen length* (see Fig. 2 in Tobias et al., 2022) combined with four categorical traits: *foraging method* (predominantly understory gleaner, ground gleaner, canopy gleaner), *diet* (insectivores or mixed diet), *nest type* (open in shrub, open on ground, cavity or open in tree) and *migration* (short migration, long migration or resident).

We derived acoustic traits calculations from the work of Krishnan & Tamma (2016). We first extracted five pure recordings without sonic background for each of the 25 candidate insectivore species from the online database Xeno-canto.org (Vellinga & Planque, 2015). We then calculated the *number of peaks* (i.e., NPIC) in the audio signal (see § Acoustic diversity, above) as well as the *frequency of the maximum amplitude peaks* for each vocal element using the “seewave” library (Sueur, Aubin & Simonis, 2008) and averaged these frequencies for each species. Being based on song and call frequency and complexity, these indices inform the adaptation of the vocal repertoire of these species to their environment.

We summarized the information conveyed by the 9 traits categories into five indices representing complementary dimensions of the functional diversity (FD) of a community (Mouillot, Graham, Villéger, Mason & Bellwood, 2013): functional richness (FRic, i.e., convex hull volume of the functional trait space summarized by a principal coordinates analysis), functional evenness (FEve, i.e., minimum spanning tree measuring the regularity of trait abundance distribution within the functional space), and functional divergence (FDiv, i.e., trait abundance distribution within the functional trait space volume) (Villéger, Mason & Mouillot, 2008), as well as Rao’s quadratic entropy (RaoQ, i.e., species dispersion from the functional centroïd) (Botta-Dukát, 2005). These were calculated for each site with the “dbFD” function of the “FD” library v.1.0.12 (Laliberté, Legendre & Shipley, 2014) in the R environment.

### Environmental data

Environmental data refer to local temperature and forest cover. We used the high 10-m resolution GIS layers from the Copernicus open platform (Cover, 2018) to calculate forest cover for all European sites. We manually calculated the percentage of forest cover for the two sites located outside Europe using the “World imagery” layer of Arcgis ver. 10.2.3552. We calculated both the percentage of forest cover in a 20-m (henceforth called *local* forest cover) and 200-m (*landscape* forest cover) buffer around the sampled oaks. We chose two nested buffer sizes to better capture the complexity of habitat structure on the diversity and acoustic activity of birds. Local forest cover is particularly important for estimating bird occurrence probability (Melles, Glenn & Martin, 2003), whereas landscape forest cover is an important predictor of bird community composition in urban areas (Rega-Brodsky & Nilon, 2017). Moreover, both local and landscape habitat factors shape insect prey distribution (Barr, van Dijk, Hylander & Tack, 2021). Preliminary analyses revealed that results were qualitatively the same using 10-, 20-or 50-m buffers as predictors of local forest cover and 200-or 500-m buffers as predictors of landscape forest cover (see Table S4.5 in Appendix S4). Because other variables were defined at the site level, we averaged the percentage of forest cover for the sampled trees per site and per buffer size.

We extracted the mean annual temperature at each site from the WorldClim database (the spatial resolution is ∼86 km^2^, Hijmans et al., 2005).

### Statistical analyses

We analyzed 14 response variables in separate linear models (LMs) (Table S2.2 in Appendix S2): bird predation attempts, species richness of the entire bird community and that of candidate insectivores, functional diversity (each of the four indices) and acoustic diversity (each of the three indices). For each response variable, we first built a full model including variables reflecting two components of the environment: climate and local habitat. The general model equation was (Eq. 1):

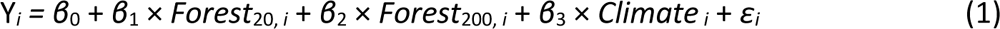

where *Y* is the response variable, *β_0_* the model intercept, *β_is_* model coefficient parameters, *Forest_20_* and *Forest_200_* the effects of the local and landscape forest cover respectively, *Climate* the effect of mean annual temperature and *ε* the residuals.

When modeling the response of bird predation attempts (Eq. 2), we added two more variables to the model, being any of the three acoustic diversity indices (*Acoustic diversity*, Eq. 2) and the species richness or any of the four indices describing the functional diversity of candidate insectivores (*Bird diversity*, Eq. 2):

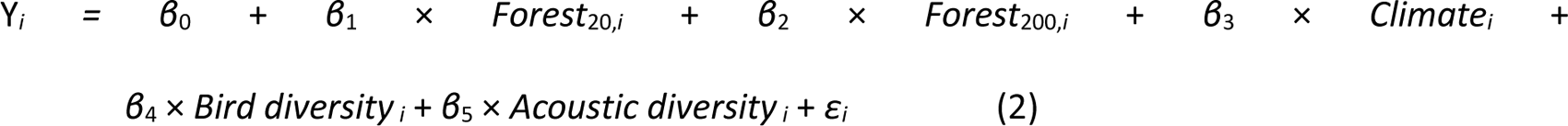

It has to be noted that the inclusion of the acoustic component in the second set of models does not imply any direct link between avian predation and acoustic diversity. By comparing models including the acoustic diversity or not, we are asking whether residual variance can be explained by this component while controlling for other sources of variation. If so, then acoustic diversity components with non-null coefficients have to be considered as proxies of predation, i.e., relatively easily measurable variables representative of unmeasured (or unknown) variables with a direct effect on predation.

We used logarithmic transformations (for bird predation attempts, acoustic entropy (H) and acoustic diversity (ADI) models) or square rooted transformation (for species richness of the complete bird community) of some response variables where appropriate to satisfy model assumptions. We scaled and centered every continuous predictor prior to modeling to facilitate comparisons of their effect sizes, and made sure that none of the explanatory variables were strongly correlated using the variance inflation factor (VIF) (all VIFs < 5, the usual cutoff values used to check for multicollinearity issues (Miles, 2014)).

For each response variable, we ran the full model as well as every model nested within the full model and then used Akaike’s Information Criterion corrected for small sample size (AICc) to identify the most effective model(s) fitting the data the best. We simultaneously selected the best variable describing the diversity and acoustic component (variable selection) and the best set of variables describing the variability of the response variable (model selection).

First, we ranked each model according to the difference in AICc between the given model and the model with the lowest AICc (ΔAICc). Models within 2 ΔAICc units of the best model (i.e., the model with the lowest AICc) are generally considered as likely (Burnham & Anderson, 2002). We computed

AICc weights for each model (*w_i_*). *w_i_* is interpreted as the probability of a given model being the best model among the set of candidate models. Eventually, we calculated the relative variable importance (RVI) as the sum of *w_i_* of every model including this variable, which corresponds to the probability a variable is included in the best model.

When several models competed with the best model (i.e., when multiple models were such that their ΔAICc < 2), we applied a procedure of multimodel inference, building a consensus model including the variables in the set of best models. We then averaged their effect sizes across all the models in the set of best models, using the variable weight as a weighting parameter (i.e., model averaging). We considered that a given predictor had a statistically significant effect on the response variable when its confidence interval excluded zero.

We run all analyses in the R language environment (R Core Team, 2020) with libraries “MuMIn” v.1.43.17 (Bartoń, 2020), “lme4” v. 1.1.27.1 (Bates, Mächler, Bolker & Walker, 2015). All R codes are provided in Appendix S5 in Supporting Information.

## Results

### Bird acoustic diversity

Of the three acoustic diversity indices (see Fig. S6.6 in Appendix S6 for correlation between indices), only Acoustic Diversity Index (ADI) and acoustic entropy (H) were significantly associated with any of the predictors tested, i.e., temperature, local forest cover and landscape forest cover (Table S2.2 in Appendix S2). ADI and H both increased with local forest cover (i.e., percentage of forest cover in a 20-m buffer around recorders). Landscape-scale forest cover (i.e., percentage of forest cover in a 200-m buffer around recorders) was the only other predictor retained in the set of competing models in a range of ΔAICc < 2 to explain acoustic entropy variation, but this predictor had little importance (RVI < 0.5) and its effect was not statistically significant (Fig. 5b; Table S2.2 in Appendix S2).

### Bird species richness and functional diversity

We identified a total of 87 bird species, among which 25 were classified as candidate functional insectivores. Bird species richness varied from 8 to 23 species per recording site (mean ± SD: 15.2 ± 3.7, n = 47 sites) and richness of candidate insectivores from 2 to 9 species (5.7 ± 1.5). The null model was among models competing in a range of ΔAICc < 2 for both total species richness and candidate insectivores (Table S2.2 in Appendix S2).

Among the five bird functional diversity and species richness indices, only functional quadratic entropy (Rao’s Q) characterizing species dispersion from the functional centroid was significantly influenced by the predictors tested (temperature, local and landscape forest cover, Table S2.2 in Appendix S2). Specifically, Rao’s Q increased with increasing temperature (Fig. 4a and Fig. 5c). Other predictors retained in the set of competing models in a range of ΔAICc < 2 had little importance (RVI < 0.5) and were not significant (Fig. 5c; Table S2.2 in Appendix S2).

**Figure 4:**
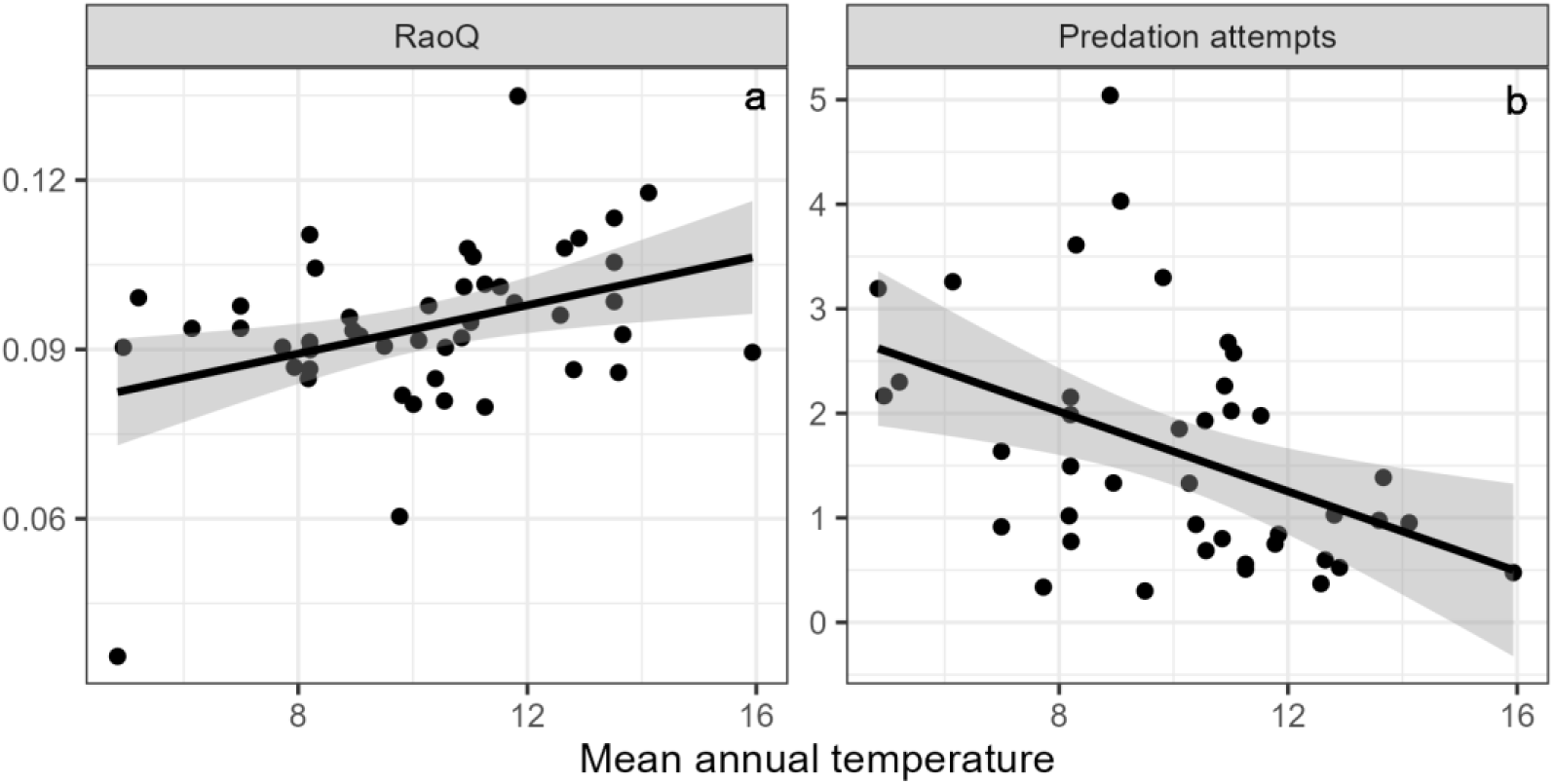
Scatter diagrams showing changes in (a) Rao’s quadratic entropy (Rao’s Q) and (b) predation attempts with mean annual temperature. These relationships were identified as significant in the linear models tested. A dot represents a site, the prediction line corresponds to a linear regression between the two variables and the gray bands represent the confidence intervals around this regression.

**Figure 5:**
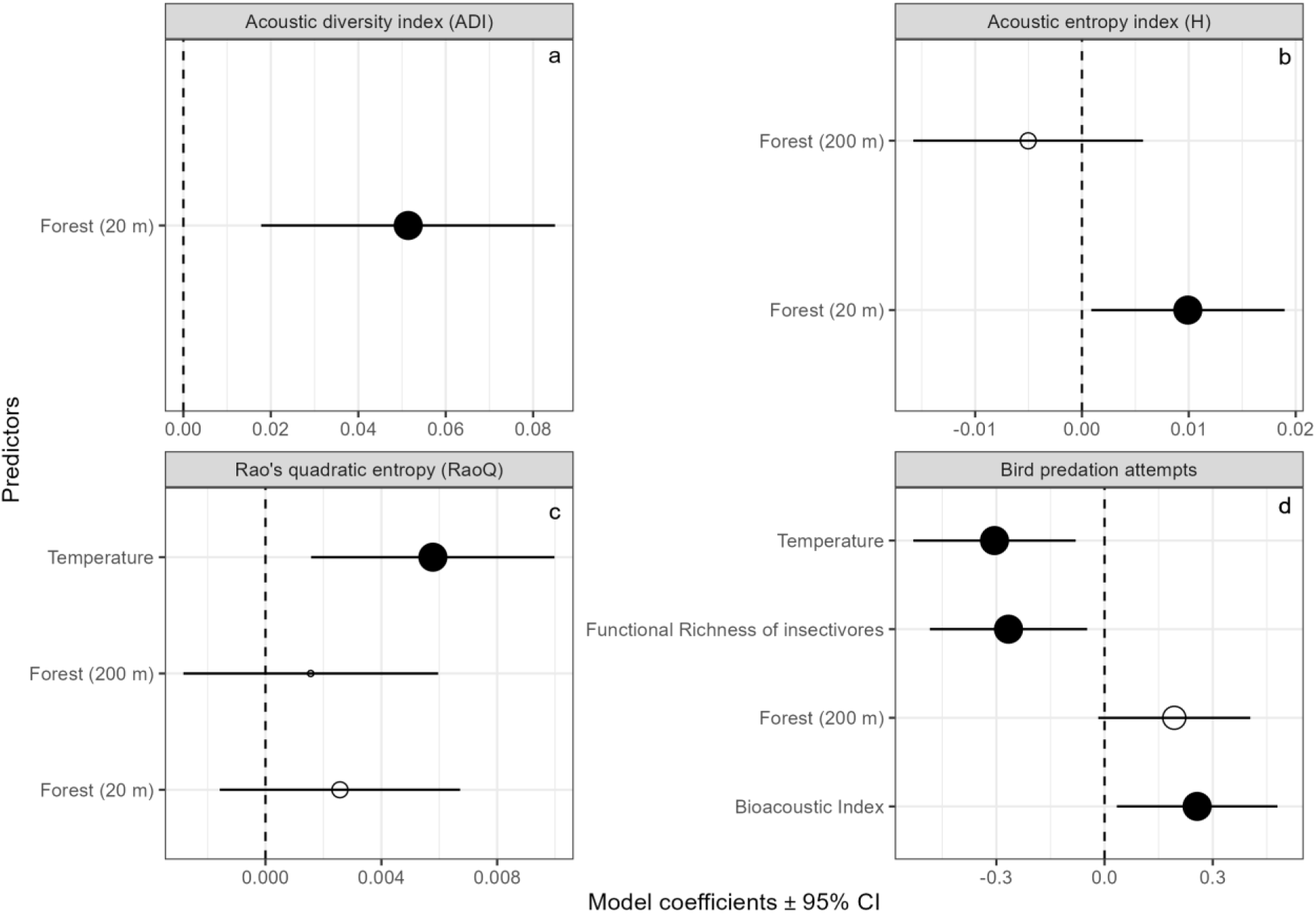
Effects of climate (described by the mean annual temperature) and habitat (percentage of forest cover at 20 or 200 m) on Acoustic Diversity Index (ADI) (a), Acoustic Entropy Index (H) (b), Rao’s quadratic entropy (RaoQ) (c), bird predation attempts (d) and effects of acoustic (Bioacoustic Index), bird diversity (Functional Richness) on bird predation attempts (d). Circles and error bars represent standardized parameter estimates and corresponding 95% confidence intervals (CI), respectively. The vertical dashed line centered on zero represents the null hypothesis. Full and empty circles represent significant and non-significant effect sizes, respectively. Circle size is proportional to RVI.

### Bird predation attempts

Of the 4,860 exposed dummy caterpillars, 22.8% (*n =* 1,108) had bird bill marks. Model selection retained two models in the set of competing models in a range of ΔAICc < 2 (Table S2.2 in Appendix S2). Bird functional richness (FRic) (RVI = 1.00), bioacoustic index (BI) (RVI = 1.00) and temperature (RVI=1.00) were selected in all models. Landscape forest cover (RVI = 0.62) was also selected in one of the two best models.

Bird predation attempts decreased with increasing mean annual temperature. Bird predation attempts further increased with bioacoustic index (BI), but decreased with bird functional richness (FRic) (Fig. 4b and Fig. 5d). This finding suggests that the acoustic component captures some features of the habitat that influence predation attempts independently of bird functional diversity. Likewise, the fact that temperature was selected a significant predictor of bird predation attempts suggests that climate has an effect on predation that is not only mediated by its effect on bird communities.

The results were comparable when we incorporated latitudinal changes in diel phenology in the calculation of predation attempts through the standardization with the daylight duration (see Table S2.2 in Appendix S2).

## Discussion

Our study confirms the well documented increase of bird diversity towards warmer regions, a pattern supporting our initial assumption that avian predation would mirror this pattern. Yet, we found the opposite – predation attempts decreased with increasing temperature – which dismissed our prediction that bird diversity and avian predation rate should correlate positively across large geographic gradients. An important result of our study is that even when the functional dimension of bird communities was accounted for, a substantial amount of variability remained to be explained and were only partially accounted for by climate-and habitat-related variables. Altogether, these findings suggest that current theory should be re-assessed, which we discuss below speculating on the main causes of deviation from theoretical expectations.

### Functional diversity of insectivorous birds and bird predation attempts are both influenced by climate, in opposite ways

In agreement with our first prediction (i, Fig. 1), we provide evidence for a significant positive relationship between temperature and the functional diversity of insectivorous birds. Despite substantial differences among functional diversity indices, this result suggests that, more functionally diverse assemblages of insectivorous birds are able to coexist locally in oak woods towards the South of Europe (Currie et al., 2004; Hillebrand, 2004; Willig et al., 2003). Of the multiple functional diversity indices commonly used to describe ecological communities, it is noticeable that only the quadratic entropy index responded positively to temperature, for it is a synthetic index that simultaneously takes into account the richness, evenness, and divergence components of functional diversity (Mouillot et al., 2013).

Contrary to our predictions (i, Fig. 1), bird predation attempts decreased with increasing temperature and were therefore inconsistently linked with bird functional diversity. More bird predation attempts at lower temperatures could be due to longer daylight duration in spring northwards, leading insectivorous birds to have more time per day to find their prey and thus allowing high coexistence of predators during a period of high resource availability (Speakman et al., 2000). Alternatively, as birds require more energy to thermoregulate in colder temperatures, they may need to feed more in order to maintain their metabolic activity (Caraco et al., 1990; Kendeigh, 1969; Steen, 1958; Wansink & Tinbergen, 1994). Moreover, temperature remained an important, significant predictor of bird predation attempts when we controlled for the duration of daylight (Table S2.2 in Appendix S2), which further supports this explanation. However, we cannot exclude the possibility that the lower predation rates at higher temperatures was due to lower prey detectability.

### Bird predation attempts are partly predicted by bird functional diversity and acoustic activity

We predicted that bird predation attempts would increase with bird abundance and functional diversity (ii, Fig. 1). The results only partially match these predictions. The relationship between bird functional diversity and predation attempts conflicted with our predictions. Specifically, we found neutral or negative relationship between these variables, depending on the functional index considered. Only functional insectivore richness was negatively correlated to predation attempts. Negative relationships between predation and predator functional diversity can arise from a combination of both intraguild predation --- predators preying upon predators (Mooney et al., 2010) - -- and intraguild competition (Houska Tahadlova et al., 2022), although we could not tease them apart in the present study. An important step forward would consist in testing whether predation patterns revealed using artificial prey are representative of predation intensity as a whole (Zvereva & Kozlov, 2021). For example, functional richness may be a proxy for dietary specialization in such a way that more functionally diverse predator communities would seek more prey of which they are specialized on and thus predate less on artificial caterpillars. It is also possible that a higher diversity of insectivorous birds in warmer regions was linked to higher diversity and abundance of arthropod prey and foraging niches (Kissling et al., 2012) and therefore to greater prey availability (Charbonnier et al., 2016). If so, then the pattern we observed may merely be representative of the ‘dilution’ of bird attacks on artificial prey among more abundant and diverse real prey (Zeuss, Brunzel & Brandl, 2017; Zvereva et al., 2019). However, the dynamics between herbivore prey abundance and predation activity are complex. A higher abundance of real herbivore prey could also lead to increased predation activity as demonstrated in studies such as Singer, Farkas, Skorik & Mooney (2011), where the presence of abundant herbivorous prey was found to drive higher predation rates by bird predators. This aligns with the notion that predator populations respond to fluctuations in prey density (Salamolard, Butet, Leroux & Bretagnolle, 2000), adjusting their foraging behavior to capitalize on available food resources. A follow-up of the present study should therefore pay special attention to the real prey density pre-existing in each sampling site where artificial prey are to be deployed as a standardized measure of predation rates across sites.

Although passive acoustic monitoring, as most other relative bird sampling methods, does not allow inferring directly bird absolute abundance, our study further brings methodological insights into the usefulness of eco-acoustics into community and functional ecology. We found that among acoustic indices that have been shown to correlate with bird abundance, activity and diversity, the Bioacoustic index was positively correlated with bird predation attempts. Yet, this index was found to be representative of the abundance and activity of singing birds (Boelman et al., 2007; Gasc et al., 2018). It is thus reasonable to infere substantial causality between vocalizing bird abundance, their acoustic activity, and the top-down control they exert upon insect prey. Such an interpretation is in line with previous studies havig reported positive relationships between bird abundance and predation attempts on artificial prey (Roels, Porter & Lindell, 2018; Sam, Koane & Novotny, 2015). It is further substantiated by the fact that if a species is recorded in a given site during the breeding season, it indicates that it is probably feeding on that territory and can potentially affect predation rates. Our study indicates that despite aknowledgeable limitations inherent to the current development of analytical tools, passive acoustic monitoring has the potential to provide acceptable proxies for the characterization of bird biodiversity, the habitat they live in, and, to some extent, the ecosystem services they provide. The present study therefore opens pathways for new research on the link between functional and acoustic ecology.

### Local forest cover predicts bird acoustic diversity, whereas landscape forest cover increases bird predation

Acoustic diversity increased with closeness of canopy cover in the immediate neighborhood (20m radius) of sampled trees (iii, Fig. 1). The most responsive indices were the Acoustic Diversity Index (ADI) and the acoustic entropy (H), both designed to predict bird acoustic diversity across different habitats under various ambient sound conditions (Fuller, Axel, Tucker & Gage, 2015; Machado, Aguiar & Jones, 2017). The former is related to a greater regularity of the soundscape and the latter is related to the amplitude between frequency bands and time. They therefore correspond to soundscapes containing multiple vocalizing species (Sueur, Pavoine, et al., 2008; Villanueva-Rivera, Pijanowski, Doucette & Pekin, 2011). Acoustic entropy is also known to respond significantly to local forest habitat (Barbaro et al., 2022), which is generally a good predictor of bird occupancy probability (Morante-Filho, Benchimol & Faria, 2021).

Bird predation attempts were best predicted by forest cover at the landscape level (Prediction (iii), Fig. 1). Indeed, it is likely that forest cover at the landscape level provides structural complexity with a dense understorey and habitat heterogeneity that is both a source of food and niches for predatory birds to exploit (Poch & Simonetti, 2013). As a result, forest cover at the landscape scale is often a key predictor of avian insectivory in various study areas (Barbaro et al., 2014; González-Gómez, Estades & Simonetti, 2006; Valdés-Correcher et al., 2021). This is also consistent with the results of Rega-Brodsky & Nilon (2017) who found greater abundance of insectivorous birds in mosaic urban or rural landscapes including a significant part of semi-natural wooded habitats, such as those we studied here.

### Large-scale variability in avian predation is not mediated by large-scale changes in bird communities

We found no evidence that the relationship between climate and bird predation attempts was mediated by changes in bird diversity or acoustic activity (iv, Fig. 1). On the contrary, climate and bird diversity and acoustic activity had independent and complementary effects on predation.

At the European scale, climate may directly drive both bird activity and abundance according to available resources (Pennings & Silliman, 2005). Even changes in the abundance of a single, particularly active, predator species along the European climatic gradient could explain the observed pattern (Maas, Tscharntke, Saleh, Dwi Putra, Clough & Siriwardena, 2015; Philpott et al., 2009). For example, the blue tit *Cyanistes caeruleus* and the great tit *Parus major* are typical and widespread canopy insectivores of European oak forests and are particularly prone to predate herbivorous caterpillars while showing considerable adaptive behavior to prey availability (Mols & Visser, 2002; Naef-Daenzer & Keller, 1999). If the predation attempts on the plasticine caterpillars were to be predominantly due to these species, then it would be their abundance and foraging activity that would play a role in predation attempts rather than the overall diversity of insectivores (Maas et al., 2015). Here, we based our assessment of functional bird composition on candidate insectivore occurrences obtained from standardized acoustic surveys, which on one hand insures that we have no observer, site, or phenological biases on species occurrences, but on the other hand also makes it difficult to precisely account for each species’ abundance. Other complementary methods to assess the relative roles of each individual bird species on predation rates should be deployed further to better account for actual predatory bird abundance and activity, including DNA sampling (Garfinkel, Minor & Whelan, 2022), camera traps (Martínez-Núñez et al., 2021) or species-specific bird surveys involving tape calls or capture methods.

## Conclusion

We found a positive association between temperature and bird functional diversity, and at the same time, a negative relationship between temperature and avian predation. Our study therefore provides partial support for the climatic clines in biodiversity hypothesis, but demonstrates that predation does not follow the same pattern. As cross-continental studies exploring the large-scale relationship between climate and the strength of biotic interactions generally ignore local factors, we argue that characterizing the contrasting habitats of the study sites is a good way to circumvent some inconsistencies in the results. We identify pre-existing real prey density and single key bird species abundances as two particularly important variables deserving further attention. Furthermore, predicting ecosystem services — here, potential pest regulation service — on a large scale by standardized proxies such as acoustic ecology for predator diversity and plasticine caterpillars for predation function seem to be good ways to reduce methodological biases and strengthen our understanding of the macro-ecology of biotic interactions.

## Supporting information

Supplementary Informations 2_3_4_6

## Acknowledgements

This study was permitted by the financial support of the BNP Paribas Foundation through its Climate & Biodiversity Initiative for the ‘Tree bodyguards’ citizen science project. FB and DT were supported by the PN 23090102 and 34PFE./30.12.2021 ‘Increasing the institutional capacity and performance of INCDS “Marin Drăcea” in the activity of RDI - CresPerfInst’ funded by the Ministry of Research, Innovation and Digitalization of Romania. MB was supported by the Forest Research Centre (CEF) (UIDB/00239/2020) and the Laboratory for Sustainable Land Use and Ecosystem Services—TERRA (LA/P/0092/2020) funded by FCT, Portugal. MdG and AK were supported by the core research group “Forest biology, ecology and technology” (P4-0107) of the Slovenian Research Agency. MVK was supported by the Academy of Finland (project 316182). VL was funded by the Czech Science Foundation (project 23-07533S) and Academy of Sciences (RVO 67985939). EM and KS were supported by the Grant Agency of the Czech Republic (19-28126X). KS and AM were supported by ERC StG BABE 805189. XM was supported by a grant from the Spanish National Research Council (2021AEP082) and a grant from the Regional Government of Galicia (IN607A 2021/03). ZV was supported by a grant awarded by the Autonomous Government of Galicia (Spain; Modalidade B-2019), and Maria Zambrano program from the Spanish Ministry of Universities. LB was supported by fundings from LTSER ZA Pyrenees Garonne. We thank the personnel of Centro Nazionale Carabinieri Biodiversità “Bosco Fontana” for help in data collection. We thank Tarja Heinonen for help in field work in Berlin, Germany. We thank Roman Hrdlička and Kari Mäntylä for bird voice identification. We thank E. L. Zvereva for help in data collection. No fieldwork permit was required for this study.

## Data availability statement

For the moment, there is an embargo on data and codes which will be lifted after open acceptance. Schille et al., 2024, « Data and codes for the article “Decomposing drivers in avian insectivory: large-scale effects of climate, habitat and bird diversity” », <u>10.57745/0E0JEA</u>, Recherche Data Gouv.

## Copyright statement

For the purpose of Open Access, a CC-BY 4.0 public copyright licence (https://creativecommons.org/licenses/by/4.0/) has been applied by the authors to the present document and will be applied to all subsequent versions up to the Author Accepted Manuscript arising from this submission.

## Biosketch

Laura Schillé is a PhD candidate interested in the functional ecology of bird communities, which she studies at different scales. She also has an interest in acoustic ecology.

Co-authors are ornithologists and/or have interests in community ecology and functional ecology.

Author contribution: B.C., L.B. and E.V.C. conceptualized the study and developed the methodology.

E.V.C., F.B., M.C.B, M.Bo., M.Br., T.D., M.dG., J.D., M.L.D., A.M.D., S.G., C.B.E., M.F., P.F.C., E.F., A.G., M.M.G., S.G., J.H., D.H., R.H., R.I., A.K., M.V.K, V.L., B.L.T., C.L.V, S.M., E.M., S.M., X.M., A.M., D.L.M., A.P., A.V.P., A.P., A.S., K.S., T.S., A.T., R.T., D.T., G.V., I.V.H., Z.V., L.B. & B.C. collected the data. A.H., M.dG., T.B., A.P., V.L., L.P., F.A., D.F., T.P., M.M., N.S., L.B., V.M., E.M., J.G., K.T., L.G., A.M., M.C.B. & R.B. processed audio recordings for bird species identification. L.S. processed and analyzed the data with guidance from B.C. and L.B. L.S., B.C., L.B. led the writing and all authors contributed critically to the revisions.

