## Supplementary Informations 2_3_4_6 for "Decomposing drivers in avian insectivory: large-scale effects of climate, habitat and bird diversity"

### **Appendix S2**

***Table S2.2*** *: Model selection table for each of the 4 types of linear models to analyze bird predation attempts (a); taxonomic diversity (b); functional diversity (c) (any of the 4 indices) and acoustic diversity (d) (any of the 3 indices). The model selection abis (in grey) corresponds to the model where the bird predation attempts variable is standardized by the daylight duration per day. The results of this model are not presented in the result part of the manuscript. Explanatory variables that cannot be included in the model selection for a given response variable are shaded. Explanatory variables included in each model are indicated by the value of the slope. Those values are in bold if the variation of the response variable as a function of the explanatory variable is significant. df is the number of parameters included in the model, logLik is the log-likelihood. Model selection is based on the Akaike information criterion adapted to small sample sizes (AICc), the difference in AICc from the best model (∆AICc) and the AICc weight of each model. The set of models with a ∆AICc less than 2 are shown. For the same response variable, if only one row is shown, it means that either the null model was selected or the best model had no competition.*

|  | **Response variable** |  | **MAT** | **Forest20** | **Forest200** | **Specific Richness** | **FRIC** | **Bioacoustic Index** | **df** | **logLik** | **AICc** | **∆AICc** | **weight** | **R^2^** |
| --- | --- | --- | --- | --- | --- | --- | --- | --- | --- | --- | --- | --- | --- | --- |
| a | Bird predation attempts |  | **-0.2808** |  | 0.1934 |  | **-0.2747** | **0.2636** | 6 | -33.722 | 82.0 | 0.00 | 0.141 | 0.354 |
|  | Bird predation attempts |  | **-0.3454** |  |  |  | **-02524** | **0.2454** | 5 | -35.615 | 83.0 | 1.01 | 0.085 | 0.300 |
| a bis | Bird predation attempts/ daylight |  | **-0.2312** |  | 0.2005 |  | **-0.2698** | **0.2603** | 6 | -33.376 | 81.3 | 0.00 | 0.118 | 0.328 |
|  | Bird predation attempts/ daylight |  | **-0.2983** |  |  |  | **-0.2466** | **0.2415** | 5 | -35.437 | 82.6 | 1.34 | 0.061 | 0.266 |
| b | Total Specific Richness |  |  |  |  |  |  |  | 2 | -31.894 | 68.1 | 0.00 | 0.388 | 0.000 |
|  | Specific Richness of Funct Insect |  |  |  |  |  |  |  | 2 | -85.026 | 174.3 | 0.00 | 0.2269 | 0.000 |
| c | FRic |  |  | 0.01329 |  |  |  |  | 3 | 76.744 | -146.9 | 0.00 | 0.328 | 0.072 |
|  | FRic |  |  |  |  |  |  |  | 2 | 74.987 | -145.7 | 1.22 | 0.178 | 0.000 |
|  | FRic |  |  | 0.02129 | -0.010510 |  |  |  | 4 | 77.238 | -145.5 | 1.42 | 0.161 | 0.089 |
|  | FEve |  |  |  |  |  |  |  | 2 | 88.033 | -171.8 | 0.00 | 0.364 | 0.000 |
|  | FDiv |  |  |  |  |  |  |  | 2 | 70.156 | -136.0 | 0.00 | 0.384 | 0.000 |
|  | RaoQ |  | **0.005503** |  |  |  |  |  | 3 | 135.765 | -265.0 | 0.00 | 0.406 | 0.138 |
|  | RaoQ |  | **0.006020** | 0.002570 |  |  |  |  | 4 | 136.583 | -264.2 | 0.76 | 0.277 | 0.164 |
|  | RaoQ |  | **0.006058** |  | 0.0015620 |  |  |  | 4 | 136.037 | -263.1 | 1.85 | 0.161 | 0.145 |
| d | BI |  | 0.5240 |  |  |  |  |  | 3 | -105.395 | 217.3 | 0.00 | 0.252 | 0.048 |
|  | BI |  |  |  |  |  |  |  | 2 | -106.581 | 217.4 | 0.09 | 0.242 | 0.000 |
|  | BI |  |  |  | -0.3827 |  |  |  | 3 | -105.956 | 218.5 | 1.12 | 0.144 | 0.026 |
|  | BI |  |  | -0.28760 |  |  |  |  | 3 | -106.230 | 219.0 | 1.67 | 0.110 | 0.015 |
|  | H |  |  | **0.008646** |  |  |  |  | 3 | 111.793 | -217.0 | 0.00 | 0.446 | 0.125 |
|  | H |  |  | **0.012540** | -0.005032 |  |  |  | 4 | 112.264 | -215.6 | 1.45 | 0.216 | 0.139 |
|  | ADI |  |  | **0.05142** |  |  |  |  | 3 | 36.744 | -66.9 | 0.00 | 0.510 | 0.171 |

### **Appendix S3**

***Table S3.3*** *: Bird species from all study sites heard in recordings and their corresponding diet in breeding season. The 25 functional insectivores are the candidates for attacking plasticine caterpillars. The comment column is used to explain why certain species were or were not considered to as functional insectivores in this study.*

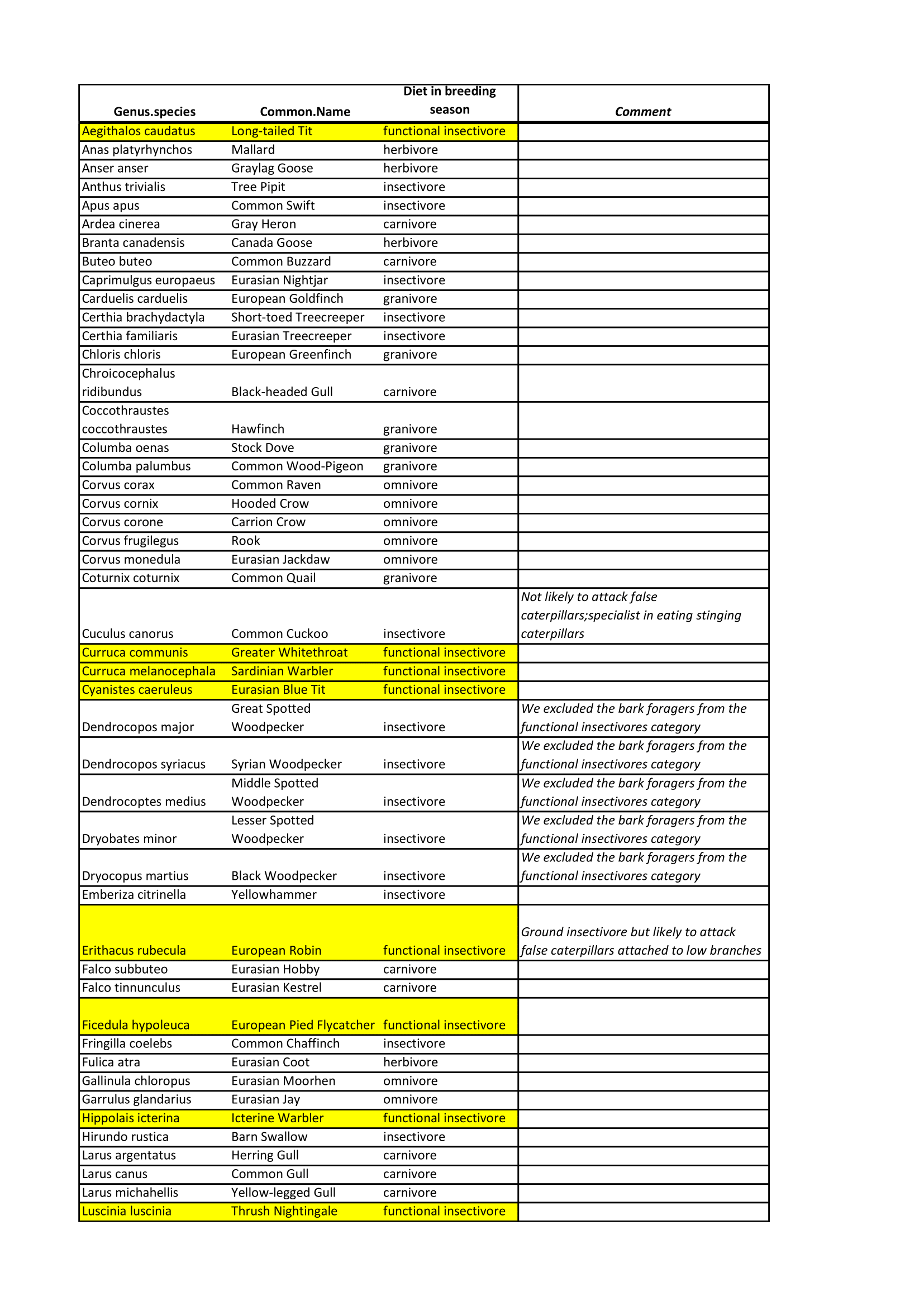

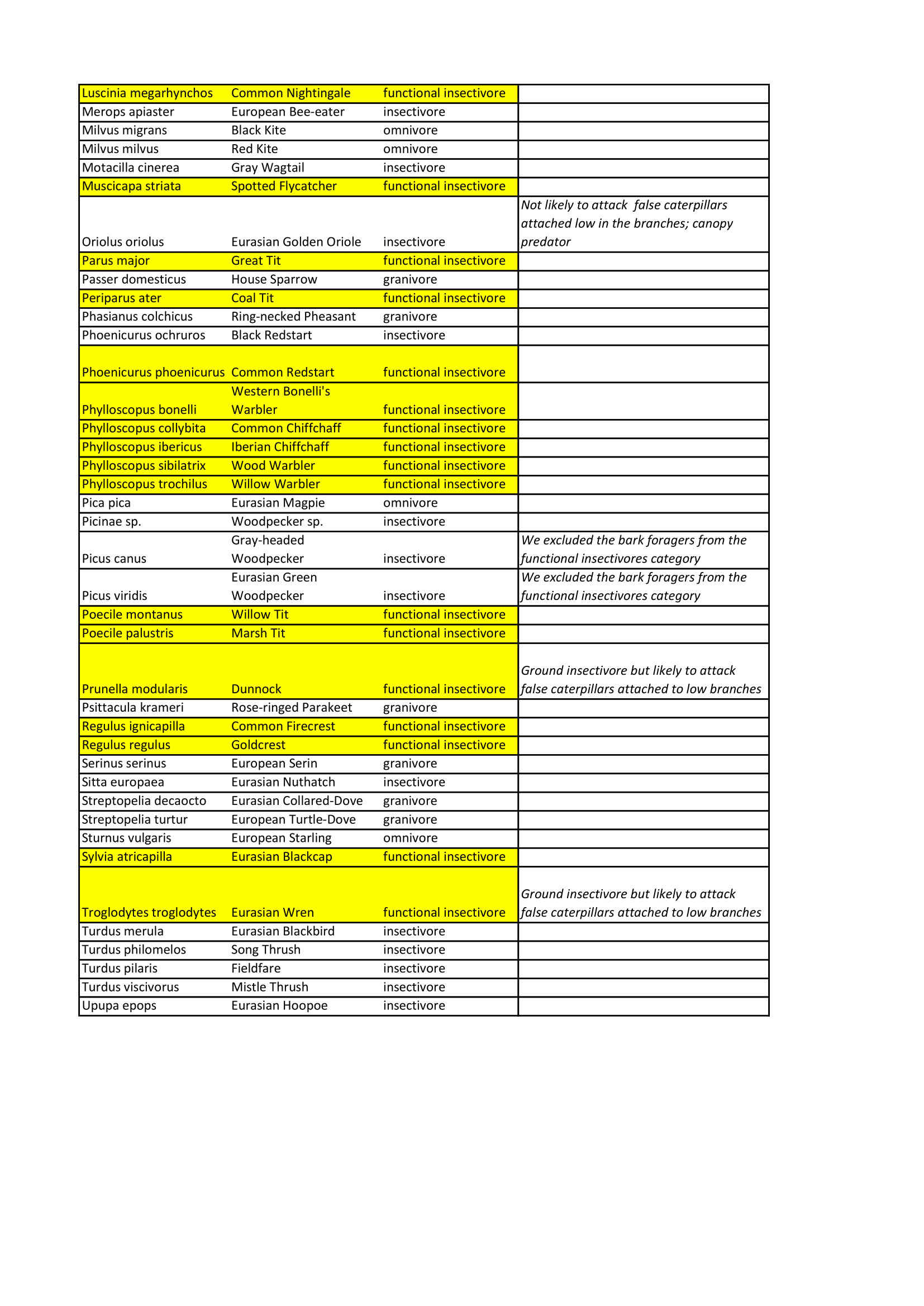

***Table S3.4*** *: Functional traits of the 25 candidate insectivores for attacking plasticine caterpillars. The colors correspond to the origin of the data specified in the legend.*

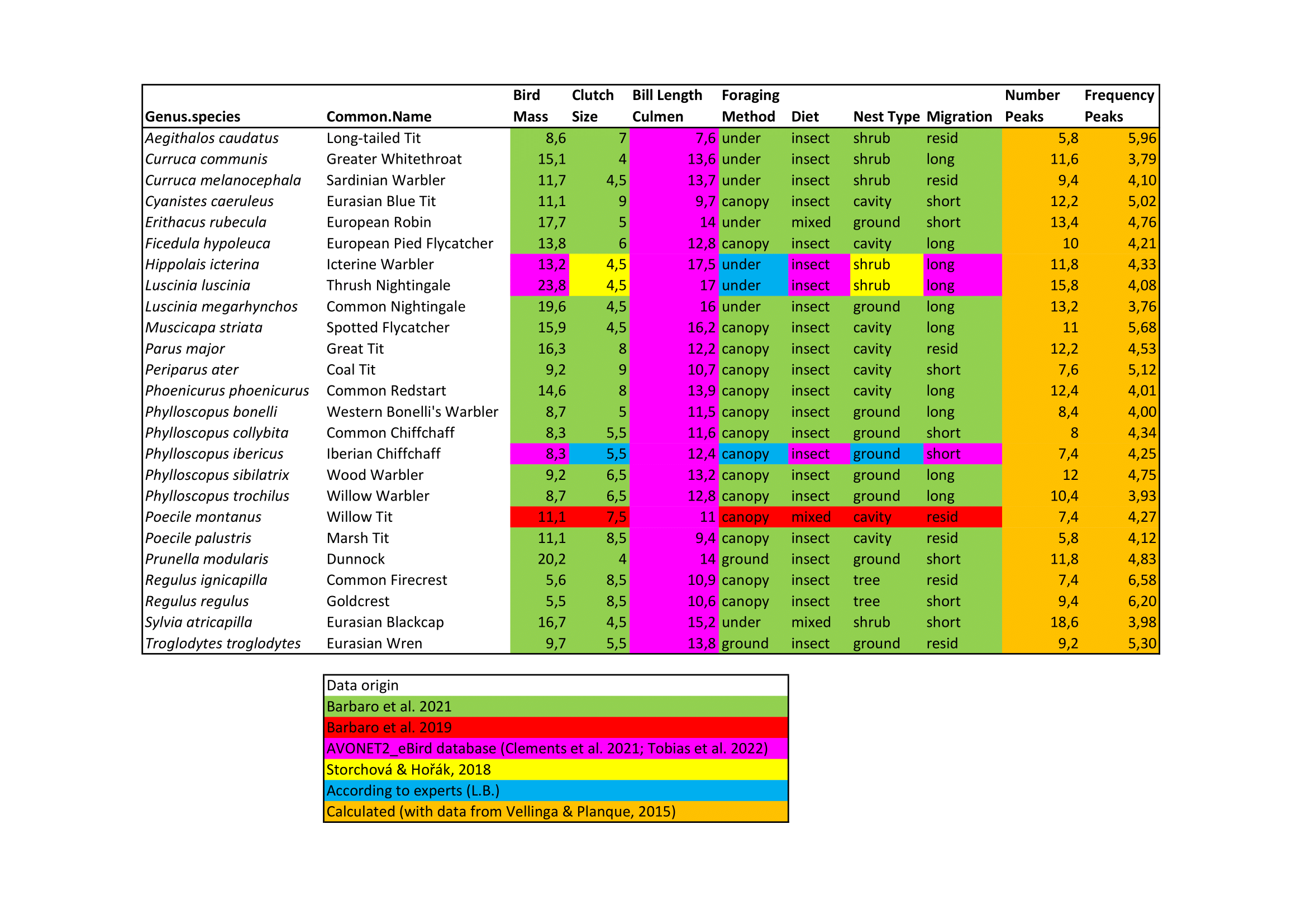

### **Appendix S4**

***Table S4.5*** *: Bird predation attempts models are presented with several combinations of predictors of forest cover at the local scale (10 m or 20 m or 50 m) and at the landscape scale (100 m or 200 m or 500 m). The + indicates a positive effect, the - a negative effect, their colour is either black for a significant effect of the predictor in the model averaging (models with deltaAICc <2) and grey for a non-significant effect of the predictor in the model averaging. We can see that the local forest cover predictors never have a significant effect on bird predation attempts. The landscape scale forest cover predictors give qualitatively identical results with buffers of 200 m and of 500 m. The buffer of 100 m caused multicollinearity issues as it was too close to the local forest cover predictors.*

| **Response variable** | **Predictors** | | | | | | |
| --- | --- | --- | --- | --- | --- | --- | --- |
|  |  | MAT | **Forest10** | **Forest100** | Specific Richness | FRIC | Bioacoustic Index |
| *Bird predation attempts* |  | - |  | + |  | - | + |
|  |  | MAT | **Forest20** | **Forest100** | Specific Richness | FRIC | Bioacoustic Index |
| *Bird predation attempts* | Multicollinearity issues between forest cover predictors | | | | | | |
|  |  | MAT | **Forest50** | **Forest100** | Specific Richness | FRIC | Bioacoustic Index |
| *Bird predation attempts* | Multicollinearity issues between forest cover predictors | | | | | | |
|  |  | MAT | **Forest10** | **Forest200** | Specific Richness | FRIC | Bioacoustic Index |
| *Bird predation attempts* |  | - | - | + |  | - | + |
|  |  | MAT | **Forest20** | **Forest200** | Specific Richness | FRIC | Bioacoustic Index |
| *Bird predation attempts* |  | - |  | + |  | - | + |
|  |  | MAT | **Forest50** | **Forest200** | Specific Richness | FRIC | Bioacoustic Index |
| *Bird predation attempts* |  | - |  | + |  | - | + |
|  |  | MAT | **Forest10** | **Forest500** | Specific Richness | FRIC | Bioacoustic Index |
| *Bird predation attempts* |  | - |  | + |  | - | + |
|  |  | MAT | **Forest20** | **Forest500** | Specific Richness | FRIC | Bioacoustic Index |
| *Bird predation attempts* |  | - |  | + |  | - | + |
|  |  | MAT | **Forest50** | **Forest500** | Specific Richness | FRIC | Bioacoustic Index |
| *Bird predation attempts* |  | - |  | + |  | - | + |

### **Appendix S6**

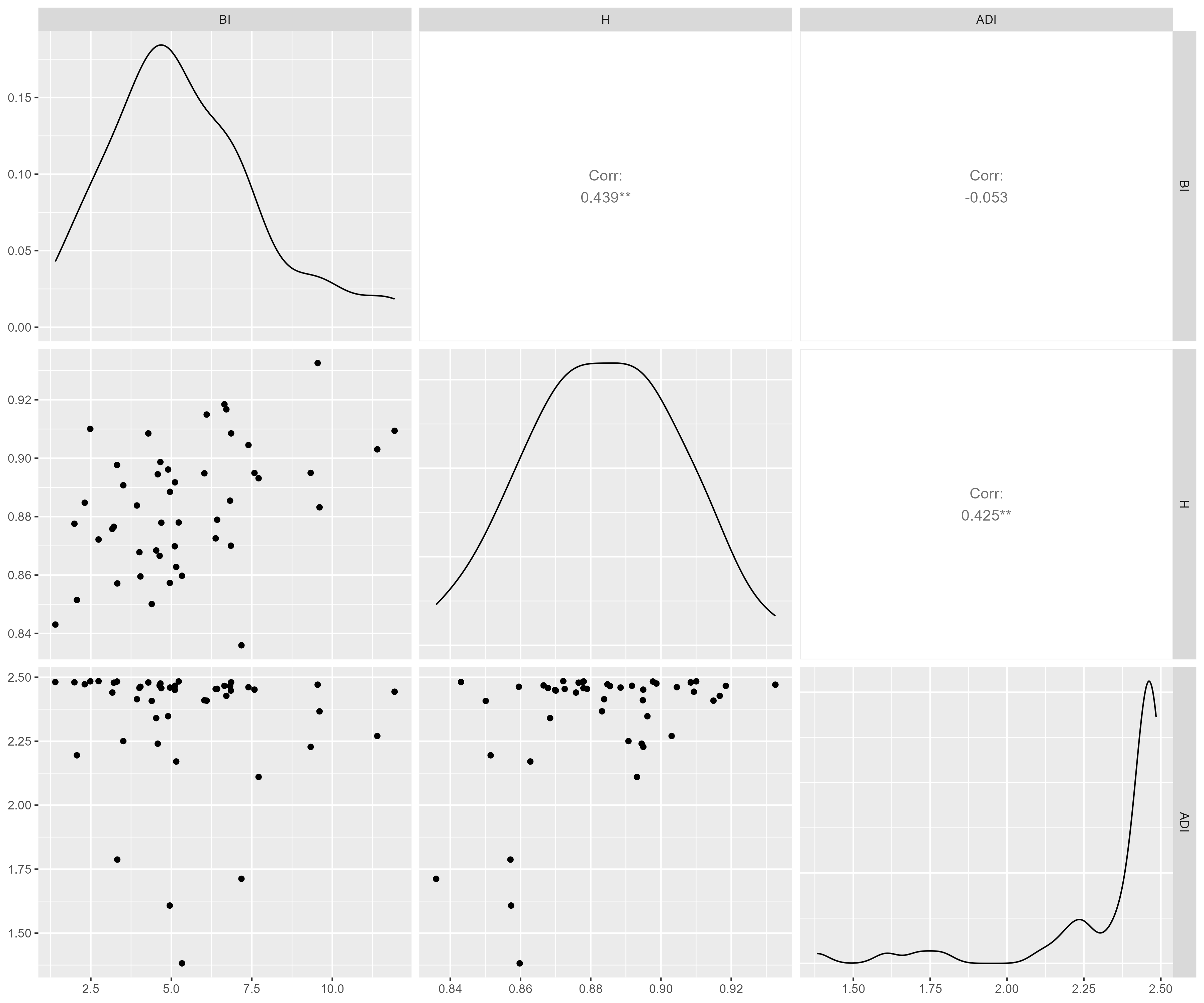

***Fig. S6.6*** *: Correlation matrix and pairwise graphical representations of acoustic diversity indices*
