## Supplementary Information 5 for "Decomposing drivers in avian insectivory: large-scale effects of climate, habitat and bird diversity"

### Latitudinal clines in bird biodiversity and predation - Rcodes

Laura Schillé

2022-08-30

```
here::i_am("Schille-et-al-2023_rev2.Rproj")
library(here)
here()
library(tidyverse)
library(ggplot2)
library(GGally)
library(ggcorrplot)
library(viridis)

library(raster)

library(sp)
library(rgeos)
library(rgdal)
library(sf)

library(doBy)

library(FD)

library(lubridate)
library(hms)

library(lme4)
```

```
library(visreg)

library(AICcmodavg)

library(MuMIn)

library(performance)

library(car)

library(DHARMa)

library(insight)


library(piecewiseSEM)
```

#### Data description

In this section, we describe the content of the archive, as well as different variables we measured. We briefly recall the methodology and indicate where the data is to be found.

The overall structure of the archive is as follows (|_ and _ refer to a folder or a file respectively; initials within brackets indicate the persons responsible for the data):

```
  |_ Schille-et-al-2023
    |_ Data
      |_ Metadata
        \_ Metadata21.csv
        \_ Metadata_acoustic_pred21.csv
      |_ Predation
        \_Data_predation21.csv
      |_ Birds
        \_ Data_bird_species21.csv
        \_ Data_bird_traits21.csv
      |_ Environement
        |_ ForestCoverCopernicus
        |_ worldclim5
        \_ Data_acoustic21.csv
        \_ Data_climate21.csv
        \_ Data_habitat21.csv
```

##### Metadata

Metadata21.csv

**Data description**

- `Site_ID`: a single identifier for each study site, there are usually 1-3 different `Tree_ID`s nested within `Site_ID`s.

- Tree_ID: a single identifier for each sampled tree.
- Year: Sampling year(2021)
- Partner_type: whether the data or biological material from which we extracted the data was a professional scientist or a teacher with her/hist pupils (Scientist, School).
- Lat and Long: the coordinates of each tree, as obtained from *google maps*.
- UTC_S_timelag: Time difference between each site and UTC
- Data_type: Are predation and acoustic data or only predation data available for each site ?

Metadata_acoustic_pred21.csv

**Data description**

- Site_ID: a single identifier for each study site, there are usually 1-3 different Tree_IDs nested within Site_IDs.
- Country
- UTC_S_timelag: Time difference between each site and UTC
- Lat and Long: the coordinates of each tree, as obtained from *google maps*.
- Record_start: Date of the first record (yyymmdd).
- Sunrise_Recordstart and Sunset_Recordstart: Time of sunrise and sunset on the date of the first record.
- Record_end: Date of the last record (yyymmdd).
- Sunrise_Recordend and Sunset_Recordend: Time of sunrise and sunset on the date of the last record.
- Week_nr: Median week number of records.
- Cater_installation: Date of installation of the false caterpillars.
- Sunrise_Instal and Sunset_Instal: Time of sunrise and sunset on the date of false caterpillars installation.
- Cater_Removal: Date of removal of the false caterpillars.
- Sunrise_Remo and Sunset_Remo: Time of sunrise and sunset on the date of false caterpillars removal.

#### Predation

Data_predation.csv

**Overview of the method** – We installed 20 dummy caterpillars made with modelling clay in oaks for 15 days, twice in a raw. We estimated predation rates as the proportion of dummy caterpillars with attack marks left by birds or arthropods. Note that organisational constraints prevented some partners to stock to the original protocol and removed caterpillars after a shorter, or longer delay.

**Data description**:

- Tree_ID: a single identifier for each sampled tree.

- `Site_ID`: a single identifier for each study site, there are usually 1-3 different `Tree_ID`s nested within `Site_ID`s.
- `Year`: Sampling year.
- `Session`: Dummy caterpillars were exposed to predators for two sessions (`Session 1`, `Session 2`) of *ca* 15 days.
- `Installation_date`: Installation date of the dummy caterpillars, for the corresponding session.
- `Removal_date`: Removal date of the dummy caterpillars, for the corresponding session.
- `N_found`: Number of dummy caterpillars found after on `Removal_date`
- `N_attacked`: Number of caterpillars with at least one predation mark, regardless of the type of mark.
- `N_bird`: Number of caterpillars with at least one predation mark left by a bird.
- `N_arthropod`: Number of caterpillars with at least one predation mark left by an arthropod.
- `N_other`: Number of caterpillars with at least one predation mark left by mammals or lezards, or unidentified predators.

#### Birds

Information on tree bird species richness and functional diversity can be extracted from the two tables we provide here.

##### Bird communities

Data_bird_species21.csv

**Overview of the method** – We identified birds from their songs, recorded through passive acoustic monitoring. There were only one recording device per site. Here, we only report a *Site* × *Species* matrix. The procedure we used to process song files is fully described in Schillé et al. (2024) **— REFERENCE TO BE ADDED SOON —**.

**Data description**:

- `Site_ID`: a single identifier for each study site, which correspond to one specific Audiomoth device.
- `Sound_sample_ID`: the identifier of the 10-minute sound sample provided to ornithologists for bird identification (4 sound samples per site).
- `Sound_sample_name`: the full name of the sound sample (of the form "SiteID_RecordingDay_RecordingHour_1st 2nd or 3rdx10minRecording.WAV").
- `Bird_species`: The latin name of bird species identified in acoustic files
- `Bird_id`: A single identifier of each bird species (according to the French Natural History Museum's Bird Banders' Code)
- `Bird_english`: The bird species English name.
- `Bird_french`: The bird species French name.

##### Bird traits

Data_bird_traits21.csv

We present here a *Species* × *Traits* matrix, aggregated from different sources (Barbaro et al. 2019; Storchová, Hořák 2018; Clements et al. 2021).

- `Bird_species`: The latin name of bird species identified in acoustic files.
- `Bird_id`: A single identifier of each bird species.
- `Forag`: Foraging method: ground gleaner (ground), understory gleaner (under), canopy gleaner (canopy), bark gleaner (bark), aerial.
- `Habitat`: Preferred habitat of the bird: forest, woodland, shrubland, grassland, rock, human modified, riverine, wetland, coastal.
- `Diet`: General diet: herb, insects, mixed, seeds, vertebrates.
- `Diet.in.breeding.season`: Diet in breeding season: herbivore, insectivore, functional insectivore, granivore, omnivore, carnivore.
- `Nest`: Nest type: open in shrub, open on ground, cavity or open in tree.
- `Migr`: Migration: short migration, long migration or resident.
- `Mass`: Bird body mass (grams).
- `Eggs`: Mean clutch size.
- `Bill.culmen.length`: Bill culmen length (mm)
- `NPIC`: Average number of peaks in the vocal phrases of functional insectivorous birds.
- `MeanFreq`: Mean frequency of the maximum amplitude peaks for each vocal phrases (kHz)

#### Environmental data

##### Acoustic

Data_acoustic21.csv

**Overview of the method** – We described the acoustic diversity of study sites using sound recorded for 30 min every 30 min for seven consecutive days between May and June 2021. Acoustic diversity indices were calculated for 1min sound samples. We calculated the median of each acoustic diversity index for a 24h period (between 00:00 and 23:59) and averaged the median values across days. The procedure we used to process acoustic data is fully described in Schillé et al. (2023)

**Data description**:

- `Site_ID`: a single identifier for each study site, which correspond to one specific Audiomoth device.
- `Record_start`: Date of the first record (yyymmdd).
- `Record_end`: Date of the last record (yyymmdd).
- `ADI`: Acoustic Diversity Index.
- `BI`: Bioacoustic Index.

- `H`: Acoustic Entropy Index.

##### Climate

Data_climate.csv

Climatic data were extracted from the WorldClim database. We provide here the codes we used for data extraction, as well as a subsamble of the data.
https://www.worldclim.org/data/worldclim21.html

**Codes** To make the code reproducible, upload the average temperature 5m data to WorldClim at this address https://www.worldclim.org/data/worldclim21.html and save it as "worldclim5". You will then be able to extract the mean annual temperature for the different sites with the commented code below. The output of the chunk will be saved and used for all further analysis.

```
#import Metadata
#Metadata21 <- read.csv2(here("Data/Metadata", "Metadata21.csv"))

#download and import worldclim data
#wd.worldclim = here("Data/Environment/worldclim5")

#Codes
#Clim <- Metadata21 %>% dplyr::select(Site_ID, Tree_ID, Lat, Long) %>%
drop_na(Lat, Long)
#tavg = raster::getData(name = "worldclim", download = T, path =
wd.worldclim, var = "tmean", res = 5, lon =  Clim$Long, lat = Clim$Lat)

#l = list(Tavg = tavg)
#wc = lapply(l, function(x) {
  #sp = SpatialPoints(coord = data.frame(x = Clim$Long, y = Clim$Lat),
proj4string = x@crs)
  #clim = rowMeans(data.frame(raster::extract(x, sp))[, 1:12])
  #return(clim)
  #})

#Clim$MAT = wc$Tavg /10

#The Clim dataframe is then saved as Data_climate.csv
```

###### Data description

- `Site_ID`: a single identifier for each for each study site.
- `Lat` and `Long`: the coordinates of each tree, as obtained from *google maps*.
- `MAT`: Mean Annual Temperature (°C ÷ 10)

##### Habitat

**Overview of the method** – We used the coordinates of oak trees to calculate the percentage of canopy cover in a 20m (local forest cover) and in a 200m (landscape forest cover) radius buffer centered upon oak trees. We used the High Resolution Layers of the CORINE land cover datasets with 10-m resolution and with reference year 2018 (± 1 year, see **Codes**, below). We assumed that landscape characteristics did not change during the survey period (2018–2020).The Russian sites were not covered by the CORINE land cover dataset. We manually estimated the percentage of impervious surfaces and forest cover using aerial images and Arcgis software.

**Codes** To make the code reproducible, upload the forest cover data to Copernicus at this address https://land.copernicus.eu/pan-european (file named: "TCD_2018_010m_eu_03035_V2_0.tif") and save it as "foresteurope_2018". You will then be able to extract the forest cover for the different buffers used with the commented code below. The output of the chunk will be saved and used for all further analysis.

```
#import Metadata
#Metadata21 <- read.csv2(here("Data/Metadata", "Metadata21.csv"))

#download and import Copernicus data
#foresteurope_2018 <-
raster(here("Data/Environment/ForestCoverCopernicus/TCD_2018_010m_eu_03035_V2
_0.tif"))

#Codes
#Habitat <- Metadata21 %>% dplyr::select(Site_ID, Tree_ID, Lat, Long) %>%
drop_na(Lat, Long)
#coordinates(Habitat) <- ~ Long + Lat

  #Buffer creation
    #20m
#Buffer20m <- gBuffer(Habitat, width = 0.0002, byid = T)
#proj4string(Buffer20m) <- CRS("+init=epsg:4326")
#CRS.new <-CRS("+init=epsg:3035")
#Buffer20m.3035 <- spTransform(Buffer20m, CRS.new)
    #200m
#Buffer200m <- gBuffer(Habitat, width = 0.002, byid = T)
#proj4string(Buffer200m) <- CRS("+init=epsg:4326")
#CRS.new <-CRS("+init=epsg:3035")
#Buffer200m.3035 <- spTransform(Buffer200m, CRS.new)

  #Extraction of forest area in 20m buffer
#For20m <- raster::extract(foresteurope_2018, Buffer20m.3035, df = T)
#For20m$For20<- For20m$TCD_2018_010m_eu_03035_V2_0
      #If 255 (outside area), NA:
#For20m <-For20m %>% naniar::replace_with_na(replace = list(For20 = 255))
#For20m<-summaryBy(For20~ID,data= For20m,FUN=mean,keep.names=T)

  #Extraction of forest area in 200m buffer
#For200m <- raster::extract(foresteurope_2018, Buffer200m.3035, df = T)
#For200m$For200<- For200m$TCD_2018_010m_eu_03035_V2_0
      # If 255, NA:
```

```
#For200m <-For200m %>% naniar::replace_with_na(replace = list(For200 = 255))
#For200m<-summaryBy(For200~ID,data= For200m,FUN=mean,keep.names=T)

  #Data aggregation
#For <- left_join(For20m, For200m, by="ID")
#Habitat <- Metadata21 %>% dplyr::select(Site_ID, Tree_ID, Lat, Long) %>%
drop_na(Lat, Long) %>% mutate(ID=rownames(Habitat))
#Habitat$ID <- as.numeric(Habitat$ID)
#Habitat <- left_join(Habitat, For, by="ID")

#The Habitat dataframe is then saved as Data_habitat21.csv
```

**Data description**

- `Site_ID`: a single identifier for each for each study site.
- `Lat` and `Long`: the coordinates of each tree, as obtained from *google maps*.
- `For20`: % forest cover in a buffer of 20m radius centered upon `Tree_ID` and averaged at the `Site_ID` level.
- `For200`: % forest cover in a buffer of 200m radius centered upon `Tree_ID` and averaged at the `Site_ID` level.

#### Data agregation

We start loading the data:

```
Metadata21 <- read.csv2(here("Data/Metadata", "Metadata21.csv"))
MetadataAcouPred <- read.csv2(here("Data/Metadata",
"Metadata_acoustic_pred21.csv"))

Climate <- read.csv2(here("Data/Environment", "Data_climate21.csv"))
Habitat <- read.csv2(here("Data/Environment", "Data_habitat21.csv"))
Acoustic <- read.csv2(here("Data/Environment", "Data_acoustic21.csv"))
Acoustic <- Acoustic[,c(1, 4:12)]

BirdSpecies <- read.csv2(here("Data/Birds", "Data_bird_species21.csv"))
BirdTraits <- read.csv2(here("Data/Birds", "Data_bird_traits21.csv"))

Pred <- read.csv2(here("Data/Predation", "Data_predation21.csv"))
```

We prepare a file with species richness and all functional diversity indices of functional insectivorous birds for each site.

```
#only one species is kept per site
BirdSpecies2 <- BirdSpecies %>%
               dplyr::select(Site_ID, Sound_sample_name, Bird_species) %>%
               group_by(Site_ID)%>%
               summarise(Bird_species=unique(Bird_species)) %>%
               mutate(Prsce=1)
```

```r
#Species richness of all species
Birds_Div <- BirdSpecies2 %>%
                group_by(Site_ID) %>%
                summarise(SpeRic_birds = n())

#diet is added to calculate an insectivore species richness
BirdTraits2 <- BirdTraits %>% dplyr::select(Bird_species,
Diet.in.breeding.season)
BirdSpecies2 <- left_join(BirdSpecies2, BirdTraits2, by="Bird_species")

Insect_Div <- BirdSpecies2 %>%
              filter(Diet.in.breeding.season=="functional insectivore") %>%
              group_by(Site_ID) %>%
              summarise(SpeRic_FunctIns = n())

Birds_Div <- left_join(Birds_Div, Insect_Div, by="Site_ID")

#### functional diversity is calculated
    #Creation of a list of functional insectivorous species
FunctInsect <- BirdSpecies2 %>%
              filter(Diet.in.breeding.season=="functional insectivore")
FunctInsect <- as.data.frame(FunctInsect)
FunctInsect <- FunctInsect %>%
               dplyr::select(Bird_species)
FunctInsect  <- unique(FunctInsect)

    #only the traits we have chosen to use are kept
BirdTraits3 <- BirdTraits %>%
               dplyr::select(Bird_species, Forag, Diet, Nest, Migr, Mass,
Eggs, Bill.culmen.length, NPIC, MeanFreq)
SpeciesTraits <- left_join(FunctInsect, BirdTraits3, by="Bird_species")
SpeciesTraits <- SpeciesTraits %>% arrange(Bird_species)

    # Traits matrix
Traits <- SpeciesTraits[-1]
rownames(Traits) <- SpeciesTraits[,1]
Traits$Forag <- as.factor(Traits$Forag)
Traits$Diet <- as.factor(Traits$Diet)
Traits$Nest <- as.factor(Traits$Nest)
Traits$Migr <- as.factor(Traits$Migr)

   # Matrix with insectivorous birds communities in rows and species in
columns
Commu.sp <- BirdSpecies2 %>%
  filter(Diet.in.breeding.season=="functional insectivore")%>%
  dplyr::select(Site_ID, Bird_species, Prsce) %>%
  pivot_wider(names_from =Bird_species, values_from = Prsce)
Commu.sp[is.na(Commu.sp)] <- 0
Commu.sp <- as.data.frame(Commu.sp)
```

```
Commu.Sp <- Commu.sp[-1]
rownames(Commu.Sp) <- Commu.sp[,1]
Commu.Sp <- as.matrix(Commu.Sp)
Commu.Sp <- Commu.Sp[,order(colnames(Commu.Sp))]

    # Distance-Based Functional Diversity Indices
DBFD <- dbFD(Traits, Commu.Sp)

DBFD

dbfd <- as.data.frame(cbind(FRic=DBFD$FRic, FEve=DBFD$FEve, FDiv=DBFD$FDiv,
RaoQ=DBFD$RaoQ))
dbfd$Site_ID <- rownames(dbfd)
dbfd <- dbfd[,c(5,1:4)]

#global dataframe with bird diversity
Birds_Div <- left_join(Birds_Div, dbfd, by="Site_ID")
```

Then, we prepare the pred file, calculating predation rates (`Predation`) as the proportion of dummy caterpillars with at least one attack mark, divided by the number of days it was exposed to predators in the field. We do it for each year and session, and then average the data at tree level, and then at site level. Because bird species were only identified in 2021, we filter the data accordingly (giving `pred_21`):

```
Pred$Installation_date <- dmy(Pred$Installation_date)
Pred$Removal_date <- dmy(Pred$Removal_date)

#Number of days between installation and removal
Pred$Nb_days <- difftime(Pred$Removal_date, Pred$Installation_date, units =
"days")
Pred <- separate (Pred, Nb_days, into=c("Nb_days",NA), sep=" ",remove=T)

Pred$Nb_days <- as.numeric(Pred$Nb_days)

#The daylight duration is added
Daylight <- MetadataAcouPred %>%
  dplyr::select(Site_ID, Sunrise_Instal, Sunset_Instal, Sunrise_Remo,
Sunset_Remo)
    #daylight duration at the installation
Daylight$Min_instal<-
period_to_seconds(as.period(as.duration(hm(Daylight$Sunset_Instal))) -
as.period(as.duration(hm(Daylight$Sunrise_Instal))))/60
    #daylight duration at the removal
Daylight$Min_remo<-
period_to_seconds(as.period(as.duration(hm(Daylight$Sunset_Remo))) -
as.period(as.duration(hm(Daylight$Sunrise_Remo))))/60
Daylight$Daylight_mean <- (Daylight$Min_instal+Daylight$Min_remo)/2
Daylight <- Daylight %>% dplyr::select(Site_ID, Daylight_mean)
Pred <- left_join(Pred, Daylight, by="Site_ID")
```

```
# Bird predation attempts are calculated through two different indices Pred_Day and Pred_Daymin:  Pred_Day = p / d, where p is the proportion of plasticine caterpillars with at least one sign of attempted predation by birds and d is the number of days plasticine caterpillars were exposed to predators in the field. Pred_Daymin = p / (d × l),  where p is the proportion of plasticine caterpillars with at least one evidence of attempted predation by birds, d is the number of days plasticine caterpillars were exposed to predators in the field, and l the median number of minutes of sunlight per day in the study area during the survey period.
Pred<- Pred %>% mutate(Pred_Daymin = ((N_bird*100)/N_found)/(Nb_days*Daylight_mean)) %>%
                    mutate(Pred_Day= ((N_bird*100)/N_found)/(Nb_days)) %>%
                    dplyr::select(Site_ID, Tree_ID, Session, Pred_Daymin, Pred_Day) %>%
                    group_by (Site_ID, Tree_ID)%>%
                    summarise(Pred_Daymin=mean(Pred_Daymin), Pred_Day=mean(Pred_Day))%>%
                    group_by(Site_ID) %>%
                    summarise(Pred_Daymin=mean(Pred_Daymin), Pred_Day=mean(Pred_Day)) %>%
                    drop_na(Pred_Daymin)
```

We combine predation data (`Pred`) with the `Birds_Div` file, the `Acoustic` file, the `Climate` file and the `Habitat` file.

```
data <- left_join(Acoustic, Pred, by="Site_ID")
data <- left_join(data, Birds_Div, by="Site_ID")
data <- left_join(data, Climate, by="Site_ID")
data <- left_join(data, Habitat, by=c("Site_ID", "Lat", "Long"))

    ## Statistical Analyses ##
```

#### Scatter diagrams showing changes in several response variables with latitude

```
scatter <- data %>%
          dplyr::select(Site_ID, MAT, Pred_Day, RaoQ)

scatter <- scatter %>% pivot_longer(cols = "Pred_Day":"RaoQ", names_to = "Variable", values_to = "Value")

scatter$Variable <- factor(scatter$Variable, levels =  c("RaoQ","Pred_Day"))

scatter <- scatter %>%
  mutate(label=ifelse(Variable=="RaoQ","a","b")) %>%
  mutate(x=16)%>%
  mutate(y=ifelse(Variable=="RaoQ",0.135,5))

Variable.labs <- c("RaoQ", "Predation attempts")
names(Variable.labs) <- c("RaoQ", "Pred_Day")
```

```
scatter %>%
  ggplot()+
  aes(x=MAT, y=Value)+
  facet_wrap(~Variable, scales = "free",
labeller=labeller(Variable=Variable.labs))+
  geom_point()+
  geom_text(data = scatter, color="black", mapping = aes(x = x, y = y, label
= label))+
  geom_smooth(data=subset(scatter,
Variable=="RaoQ"|Variable=="Pred_Day"),method = "lm", color="black")+
  ylab("")+
  xlab("Mean annual temperature")+
  theme_bw()
```

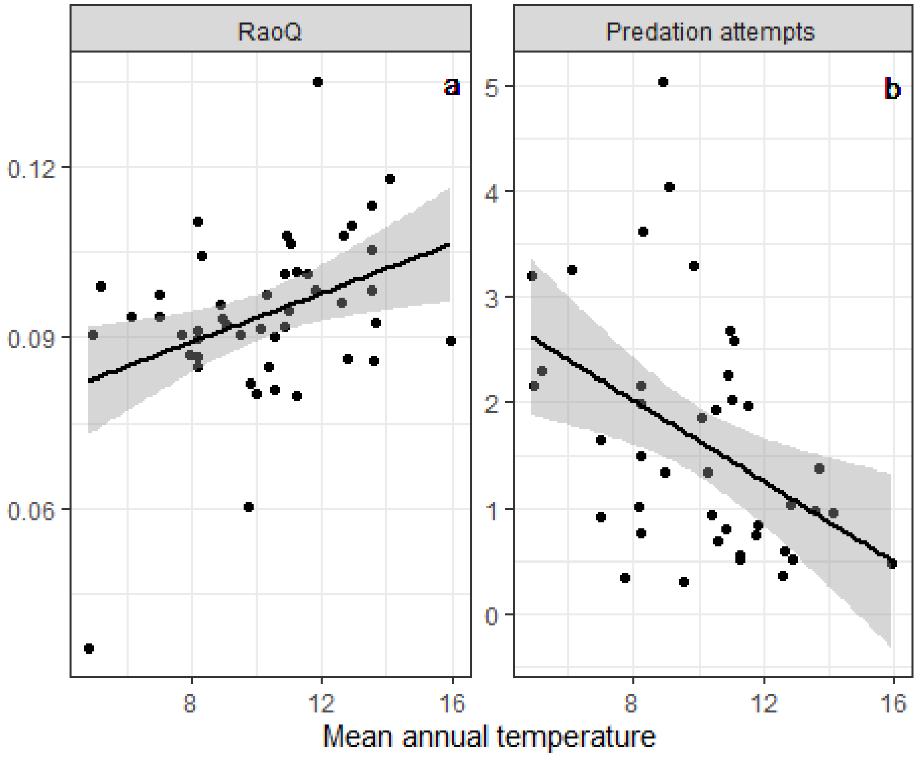


```
#ggsave(here("Figures", "ScatterLat.tiff"), width = 6, height = 3)
```

```
data2 <- data %>% dplyr::select(Site_ID, Pred_Day, Pred_Daymin, Lat, MAT,
For20, For200, SpeRic_birds, SpeRic_FunctIns, FRic , FEve , FDiv , RaoQ , BI
, H , ADI)
```

#### Statistical models

#### I- Models on predation attempts

```
## We scale the variables used in the predation models
dScale <- cbind(data2[,c(1:3)], scale(data2[4:16])) %>% drop_na()
```

#### 1- Models on predation index not standardised by daylight duration

```
hist((dScale$Pred_Day))

hist(log(dScale$Pred_Day)) #the log transformation of the variable follows a
normal distribution

mPred = lm(log(Pred_Day) ~  MAT  +For20 +For200 +  BI + H + ADI +
SpeRic_FunctIns +FRic + FEve + FDiv+ RaoQ, data = dScale, na.action =
"na.fail")

(mslPred = dredge(mPred, subset =
                #!(Lat & MAT) &
                !(SpeRic_FunctIns & FRic) &
                !(SpeRic_FunctIns & FEve) &
                !(SpeRic_FunctIns & FDiv) &
                !(SpeRic_FunctIns & RaoQ) &
                !(FRic & FEve)&
                !(FRic & FDiv)&
                !(FRic & RaoQ)&
                !(FEve & FDiv)&
                !(FEve & RaoQ)&
                !(FDiv & RaoQ)&
                ! (BI & H) &
                ! (BI & ADI) &
                ! (H & ADI),
                extra = list(R2=function(x) {r.squaredGLMM(x,
null=nullmodel)})))

vif(mPred)

top.modelsPred <- get.models(mslPred, subset=delta<2)
avgPred <- model.avg(top.modelsPred, subset=delta<2)
confint(avgPred)

coef(avgPred)

summary(avgPred)

MuMIn::importance(avgPred)
```

#### 1- Models on predation index standardised by daylight duration

```
hist((dScale$Pred_Daymin))

hist(log(dScale$Pred_Daymin)) #the log transformation of the variable follows
a normal distribution
```

```
mPred2 = lm(log(Pred_Daymin) ~  MAT + For20 +For200 +  BI + H  + ADI +
SpeRic_FunctIns  +FRic + FEve + FDiv+ RaoQ, data = dScale, na.action =
"na.fail")
```

```
(mslPred2 = dredge(mPred2, subset =
                !(SpeRic_FunctIns & FRic) &
                !(SpeRic_FunctIns & FEve) &
                !(SpeRic_FunctIns & FDiv) &
                !(SpeRic_FunctIns & RaoQ) &
                !(FRic & FEve)&
                !(FRic & FDiv)&
                !(FRic & RaoQ)&
                !(FEve & FDiv)&
                !(FEve & RaoQ)&
                !(FDiv & RaoQ)&
                ! (BI & H) &
                ! (BI & ADI) &
                ! (H & ADI),
                extra = list(R2=function(x) {r.squaredGLMM(x,
null=nullmodel)})))

vif(mPred2)

top.modelsPred2 <- get.models(mslPred2, subset=delta<2)
avgPred2 <- model.avg(top.modelsPred2, subset=delta<2)
confint(avgPred2)

coef(avgPred2)

summary(avgPred2)

MuMIn::importance(avgPred2)
```

#### II- Models on acoustic diversity

#### Correlation table between acoustic indices

```
AcouInd <- as.data.frame(data2[,c("BI","H","NPIC")])

#Function that calculates correlation matrices
panel.cor <- function(x, y, digits=2, prefix="", cex.cor) {
usr <- par("usr"); on.exit(par(usr))
par(usr = c(0, 1, 0, 1))
r <- cor(x, y)
txt <- format(c(r, 0.123456789), digits=digits)[1]
txt <- paste(prefix, txt, sep="")
if(missing(cex.cor))
 cex <- 0.8/strwidth(txt)
test <- cor.test(x,y)
Signif <- symnum(test$p.value, corr = FALSE, na = FALSE,
cutpoints = c(0, 0.001, 0.01, 0.05, 0.1, 1),
symbols = c("***", "**", "*", ".", " "))
text(0.5, 0.5, txt)
text(.8, .8, Signif, cex=cex, col=2)
}
```

```
ggpairs(AcouInd,
 lower.panel=panel.cor, upper.panel=panel.smooth, main = "Correlation between
acoustic indices")
```

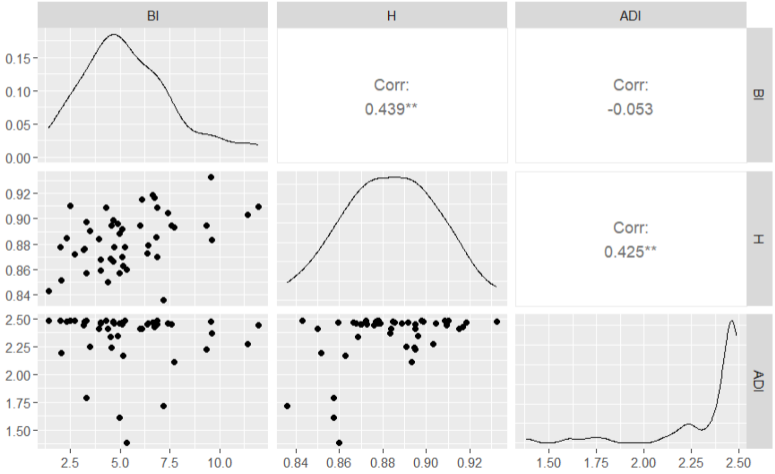


```
#ggsave(here("Figures", "AcouCorr.tiff"), width = 12, height = 10)

### Same with functional diversity indices calculated with and without bird
species occurancy
FuncDiv <-  data2 %>%
  dplyr::select(SpeRic_birds:RaoQ)

ggpairs(FuncDiv,
 lower.panel=panel.cor, upper.panel=panel.smooth, main = "Correlation between
functional diversity indices")
```

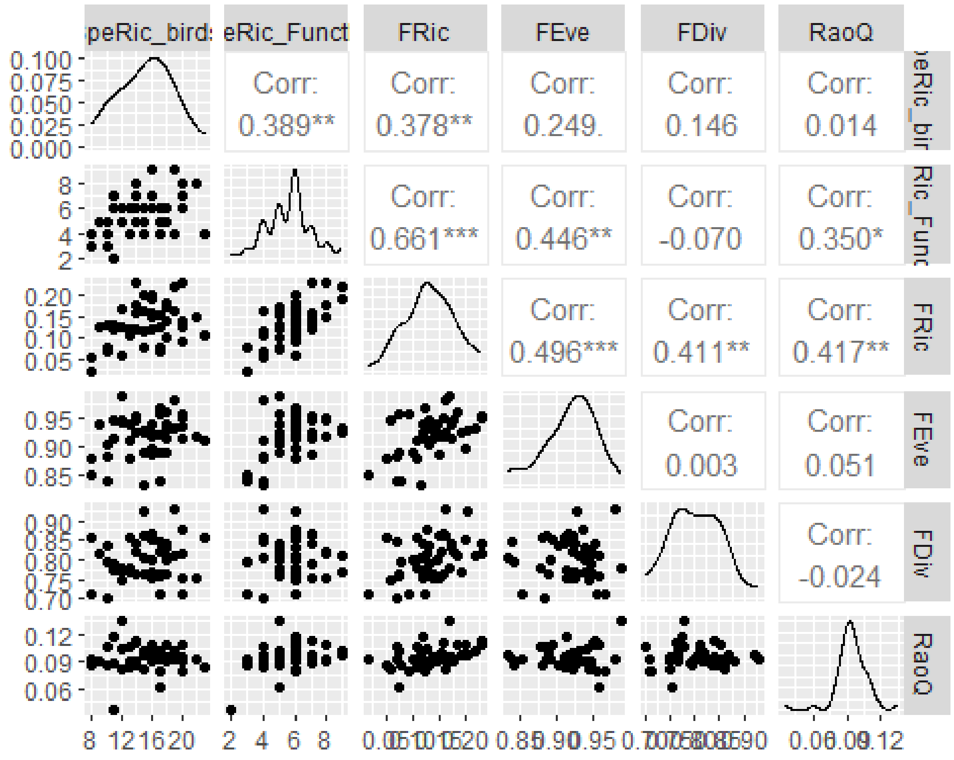


```
## We scale the variables used in the acoustic diversity  models
dScaleAcou <- cbind(data2[,c(1,14:16)], scale(data2[4:7])) %>% drop_na()
```

#### 1- Models on ADI

```
hist(dScaleAcou$ADI)

hist(log(dScaleAcou$ADI))

mH = lm(log(ADI) ~ MAT + For20 + For200, data = dScaleAcou, na.action =
"na.fail")

(mslADI = dredge(mADI, #subset = !(Lat & MAT),
extra = list(R2=function(x) {r.squaredGLMM(x, null=nullmodel)})))

vif(mADI)

top.modelsADI <- get.models(mslADI, subset=delta<2) #only one model


bestADI <- lm(log(ADI) ~ For20, data = dScaleAcou, na.action = "na.fail")

summary(bestADI)
```

#### 2- Models on H

```
hist(dScaleAcou$H)
```

```
hist(log(dScaleAcou$H)) #the log transformation of the variable follows a normal distribution

mH = lm(log(H) ~ MAT + For20 + For200, data = dScaleAcou, na.action = "na.fail")

(mslH = dredge(mH, #subset = !(Lat & MAT),
extra = list(R2=function(x) {r.squaredGLMM(x, null=nullmodel)})))

vif(mH)

top.modelsH <- get.models(mslH, subset=delta<2)
avgH <- model.avg(top.modelsH, subset=delta<2)
confint(avgH)

coef(avgH)

summary(avgH)

MuMIn::importance(avgH)
```

#### 3- Models on BI

```
mBI = lm(BI ~ MAT + For20 + For200 , data = dScaleAcou, na.action = "na.fail")

(mslBI = dredge(mBI, #subset = !(Lat & MAT),
extra = list(R2=function(x) {r.squaredGLMM(x, null=nullmodel)})))

vif(mBI)

top.modelsBI <- get.models(mslBI, subset=delta<2)
avgBI <- model.avg(top.modelsBI, subset=delta<2)
confint(avgBI)

coef(avgBI)

summary(avgBI)

MuMIn::importance(avgBI)
```

### III- Models on functional diversity

#### Correlation table between acoustic indices and functional diversity indices

```
Bird <- data2 %>% dplyr::select(SpeRic_birds, FRic:ADI) %>% drop_na()

Birdcor <- round(cor(Bird),1)

ggcorrplot(Birdcor, hc.order = TRUE, type = "lower",
     outline.col = "white", title = "Correlation matrix between acoustic indices
        and functional diversity indices")
```

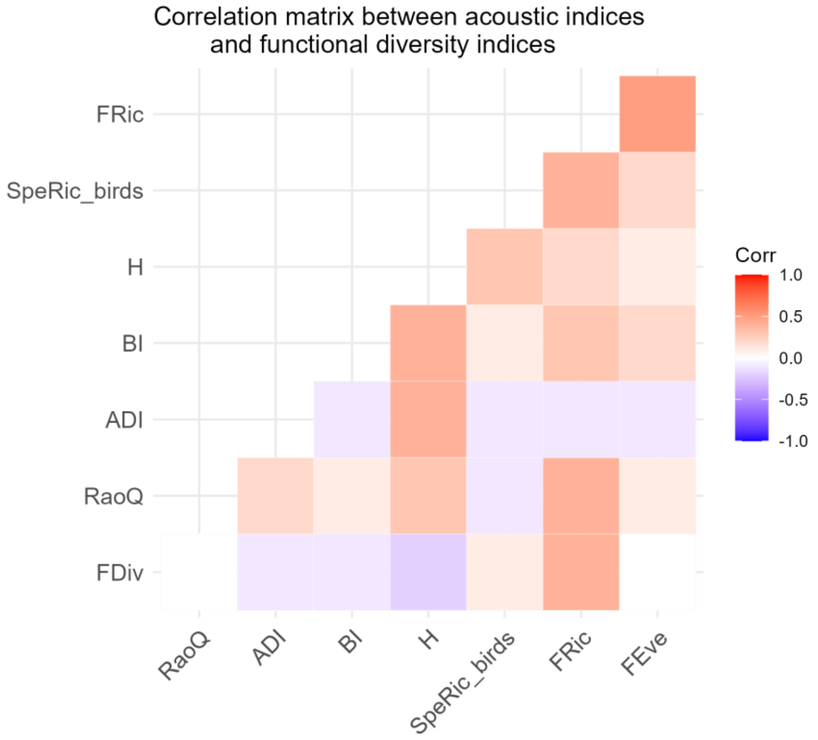


```
#ggsave(here("Figures", "AcouDivIndCorr.tiff"), width = 6, height = 6)

## We scale the variables used in the functional diversity models
dScaleRic <- cbind(data2[,c(1,8:13)], scale(data2[4:7]))
```

#### 1- Models on species richness of all birds

```
hist(dScaleRic$SpeRic_birds)

hist(sqrt(dScaleRic$SpeRic_birds)) #the sqrt transformation of the variable
follows a normal distribution
```

```
mRicTot = lm(sqrt(SpeRic_birds) ~ MAT + For20 + For200, data = dScaleRic,
na.action = "na.fail")

(mslRicTot = dredge(mRicTot, #subset = !(Lat & MAT),
extra = list(R2=function(x) {r.squaredGLMM(x, null=nullmodel)})))

vif(mRicTot)

top.modelsRicTot <- get.models(mslRicTot, subset=delta<2) ## only one model
```

#### 2- Models on species richness of functional insectivorous birds

```
hist((dScaleRic$SpeRic_FunctIns))
```

```
mRicFI = lm((SpeRic_FunctIns) ~ MAT + For20 + For200, data = dScaleRic,
na.action = "na.fail")

(mslRicFI = dredge(mRicFI, #subset = !(Lat & MAT),
extra = list(R2=function(x) {r.squaredGLMM(x, null=nullmodel)})))

vif(mRicFI)

top.modelsRicFI <- get.models(mslRicFI, subset=delta<2)
avgRicFI <- model.avg(top.modelsRicFI, subset=delta<2)
confint(avgRicFI)

coef(avgRicFI)

summary(avgRicFI)

MuMIn::importance(avgRicFI)
```

#### 3- Models on FRic

```
dScaleRic2 <- dScaleRic %>% drop_na()

hist(dScaleRic2$FRic)
```

```
mFRic = lm((FRic) ~ MAT + For20 + For200, data = dScaleRic2, na.action =
"na.fail")

(mslFRic = dredge(mFRic, #subset = !(Lat & MAT),
extra = list(R2=function(x) {r.squaredGLMM(x, null=nullmodel)})))

vif(mFRic)

top.modelsFRic <- get.models(mslFRic, subset=delta<2)
avgFRic <- model.avg(top.modelsFRic, subset=delta<2)

coef(avgFRic)

summary(avgFRic)

MuMIn::importance(avgFRic)
```

#### 4- Models on FEve

```
hist((dScaleRic2$FEve))

mFEve = lm((FEve) ~  MAT + For20 + For200, data = dScaleRic2, na.action =
"na.fail")

(mslFEve = dredge(mFEve, #subset = !(Lat & MAT),
extra = list(R2=function(x) {r.squaredGLMM(x, null=nullmodel)})))
```

```
vif(mFEve)

top.modelsFEve <- get.models(mslFEve, subset=delta<2)
avgFEve <- model.avg(top.modelsFEve, subset=delta<2)
confint(avgFEve)

coef(avgFEve)

summary(avgFEve)

MuMIn::importance(avgFEve)
```

#### 5- Models on FDiv

```
hist((dScaleRic2$FDiv))

mFDiv = lm((FDiv) ~ MAT + For20+  For200, data = dScaleRic2, na.action =
"na.fail")

(mslFDiv = dredge(mFDiv, #subset = !(Lat & MAT),
 extra = list(R2=function(x) {r.squaredGLMM(x, null=nullmodel)})))

vif(mFDiv)

top.modelsFDiv <- get.models(mslFDiv, subset=delta<2)
avgFDiv <- model.avg(top.modelsFDiv, subset=delta<2)
confint(avgFDiv)

coef(avgFDiv)

summary(avgFDiv)

MuMIn::importance(avgFDiv)
```

#### 6- Models on RaoQ

```
hist((dScaleRic$RaoQ))

mRaoQ = lm((RaoQ) ~ MAT + For20 + For200, data = dScaleRic, na.action =
"na.fail")

(mslRaoQ = dredge(mRaoQ, #subset = !(Lat & MAT),
extra = list(R2=function(x) {r.squaredGLMM(x, null=nullmodel)})))

vif(mRaoQ)

top.modelsRao <- get.models(mslRaoQ, subset=delta<2)
avgRao <- model.avg(top.modelsRao, subset=delta<2)
confint(avgRao)

coef(avgRao)

summary(avgRao)
```

```
MuMIn::importance(avgRao)
```

#### IV- Summary figure

```
my_table = function(my_model, model_lab = "Model") {

  if(length(my_model) > 1) {
    RVI = c("", round(c(importance(my_model)),2))
    RVI_names = c("(Intercept)", names(importance(my_model)))
    RVI = data.frame(Predictors = RVI_names, RVI = RVI)

    full_join(RVI,
            data.frame(Model = model_lab,
                       Predictors = names(coef(my_model)),
                       est = coef(my_model),
                       lci = confint(my_model)[,1],
                       uci = confint(my_model)[,2]),
            by = "Predictors") %>%
    mutate(s = ifelse(lci < 0 & uci > 0, "ns", "*"))

  }else{

    RVI = data.frame(Predictors = names(coef(my_model[[1]])), RVI = rep(1,
length(names(coef(my_model[[1]])))))

    full_join(RVI,
            data.frame(Model = model_lab,
                       Predictors = names(coef(my_model[[1]])),
                       est = coef(my_model[[1]]),
                       lci = confint(my_model[[1]])[,1],
                       uci = confint(my_model[[1]])[,2]),
            by = "Predictors") %>%
    mutate(s = ifelse(lci < 0 & uci > 0, "ns", "*"))
  }

}



x =
  rbind(
  my_table(avgPred, "Bird predation attempts"),
  my_table(avgH, "Acoustic entropy index (H)"),
  my_table(get.models(mslADI, subset=delta==0), "Acoustic diversity index
(ADI)"),
  my_table(avgRao, "Rao's quadratic entropy (RaoQ)")) %>%
  filter(Predictors != "(Intercept)")

x$Predictors[x$Predictors == "For200"] = "Forest (200 m)"
x$Predictors[x$Predictors == "MAT"] = "Temperature"
x$Predictors[x$Predictors == "For20"] = "Forest (20 m)"
```

```r
x$Predictors[x$Predictors == "FRic"] = "Functional Richness of insectivores"
x$Predictors[x$Predictors == "BI"] = "Bioacoustic Index"


x <- x %>%
  mutate(RVI = as.numeric(RVI)) %>%
  mutate(RVI.size=0.1*RVI)%>%
  mutate(Component = ifelse(Predictors=="Forest (200 m)", "Habitat",
ifelse(Predictors=="Forest (20 m)", "Habitat", ifelse(Predictors=="Latitude",
"Geography", ifelse(Predictors=="Temperature", "Climate",
ifelse(Predictors=="Functional Richness of insectivores", "Bird diversity",
"Acoustic"))))))

x$Model <- as.factor(x$Model)
x$group <- factor(x$Model, levels =  c("Acoustic diversity index
(ADI)","Acoustic entropy index (H)", "Rao's quadratic entropy (RaoQ)", "Bird
predation attempts" ))


## letters on facet_wrap
x$label <- c("d","d","d","d"","b","b","a","c","c","c")
x$y <- c(0.5,0.5,0.5,0.5,0.019,0.019,0.085,0.010,0.010,0.010)
x$x <- c(4.4,4.4,4.4,4.4,2.5,2.5,1.55,3.4,3.4,3.4)

 x %>% ggplot()+
  aes(Predictors, est, shape = s, group=group)+
  geom_point(aes(size=RVI))+
  guides(size=FALSE)+
  geom_hline(yintercept =  0, lty = 2) +
  geom_linerange(aes(ymin = lci, ymax = uci)) +
  geom_linerange(aes(ymin = lci, ymax = uci)) +
  facet_wrap(~group, scales = "free", ncol=, nrow=2) +
  geom_text(data = x, color="black", mapping = aes(x = x, y = y, label =
label))+
  scale_shape_manual(values = c(16, 1), guide = NULL) +
  ylab("Estimate")+
  labs(y=expression("Model coefficients" %+-% "95% CI", x=""))+
  coord_flip() +
  theme_bw()
```

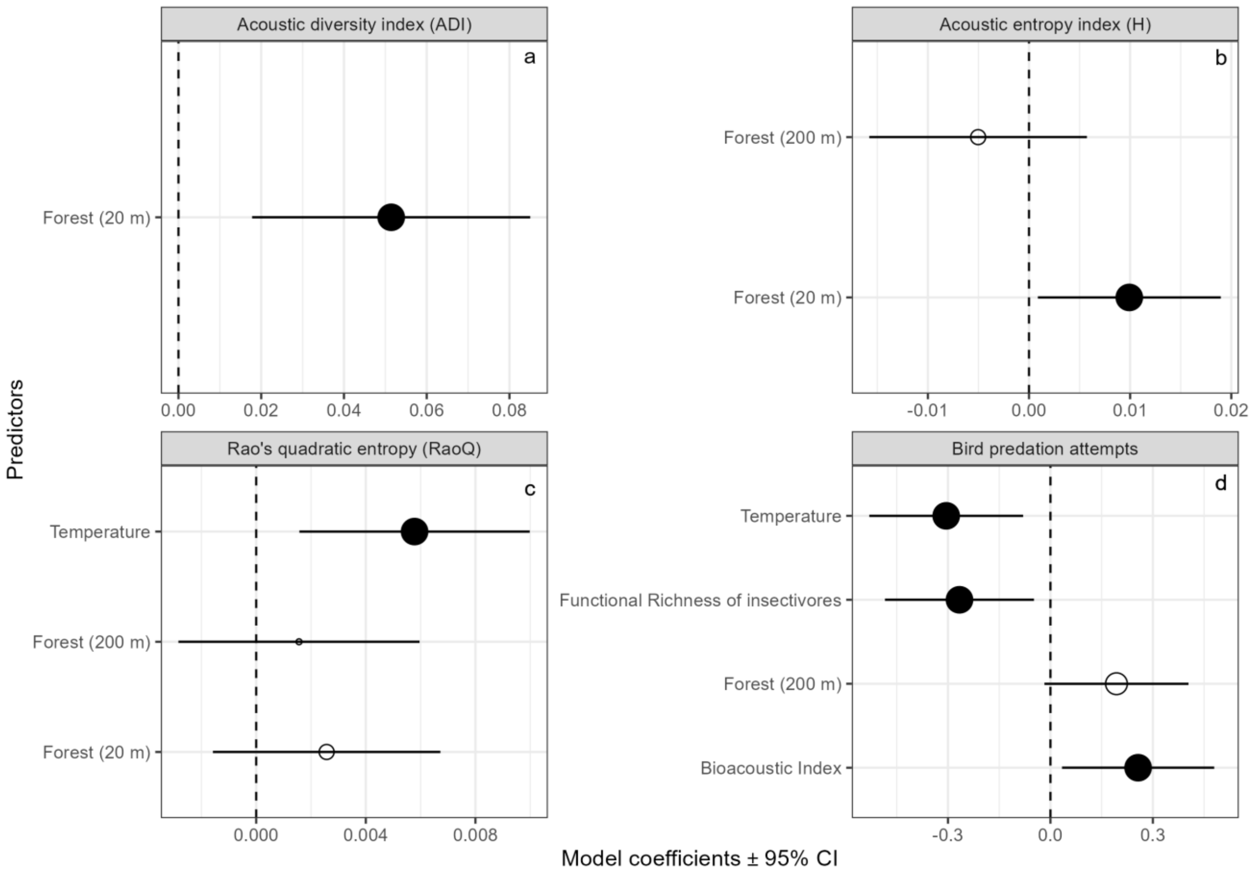


```
#ggsave(here("Figures", "GlobFig.tiff"), width = 8.5, height = 6)
```
